# Dorsal horn CGRP-expressing interneurons contribute to nerve injury-induced mechanical hypersensitivity

**DOI:** 10.1101/2020.06.08.131839

**Authors:** LS Löken, A Etlin, M Bernstein, M Steyert, J Kuhn, K Hamel, I Llewellyn-Smith, JM Braz, AI Basbaum

## Abstract

Primary sensory neurons are generally considered the only source of dorsal horn calcitonin gene-related peptide (CGRP), a neuropeptide critical to the transmission of pain messages. Using a tamoxifen-inducible CGRP^CreER^ transgenic mouse, here we identified a distinct population of CGRP-expressing excitatory interneurons in lamina III of the spinal cord dorsal horn and trigeminal nucleus caudalis. These interneurons have spine-laden, dorsally-directed, dendrites and ventrally-directed axons. Neither innocuous nor noxious stimulation provoked significant Fos expression in these neurons. However, synchronous, electrical non-nociceptive Aβ primary afferent stimulation of dorsal roots depolarized the CGRP interneurons, consistent with their receipt of a VGLUT1 innervation. In contrast, chemogenetic activation produced a significant mechanical hypersensitivity. Importantly, the CGRP interneurons could be activated after peripheral nerve injury, but only with concurrent innocuous, brush stimulation. These findings suggest that hyperexcitability of dorsal horn CGRP interneurons is an important contributor to the circuits that render touch painful after peripheral nerve damage.

## INTRODUCTION

Calcitonin Gene Related Peptide (CGRP) is the most prominent molecular marker of the peptidergic subpopulation of primary afferent nociceptors (Basbaum, Bautista, Scherrer, & Julius, 2009). When released from peripheral terminals of sensory neurons, CGRP acts on endothelial cells that line blood vessels, producing pronounced vasodilation (Brain, Williams, Tippins, Morris, & MacIntyre, 1985). Recent efforts to develop novel therapeutics in the management of migraine led to the successful development of antibodies that scavenge CGRP, reducing the vasodilation that triggers migraine (Ho, Edvinsson, & Goadsby, 2010). When released into the superficial dorsal horn from the central branches of sensory neurons, CGRP, along with its co-occurring neuropeptide, substance P, potentiates the glutamatergic excitation of postsynaptic neurons, contributing to injury-provoked central sensitization (Ryu, Gerber, Murase, & Randic, 1988; Woolf & Wiesenfeld-Hallin, 1986). The latter process, in turn, contributes to the ongoing pain and profound hypersensitivity characteristic of both inflammatory and neuropathic pains. Interestingly, a recent study showed that pharmacological inhibition of CGRP receptor signaling in the periphery alleviates incision-induced mechanical and heat hypersensitivity, but not neuropathic pain, suggesting that primary sensory neuron-derived CGRP differentially influences injury-induced persistent pain (Cowie, Moehring, O’Hara, & Stucky, 2018).

Despite a much earlier study in which colchicine was used to enhance somatic CGRP levels (Kruger, Sternini, Brecha, & Mantyh, 1988; Tie-Jun, Xu, & Hokfelt, 2001) and a more recent report (McCoy, Taylor-Blake, & Zylka, 2012) of small CGRP-positive cells in the dorsal horn of a reporter mouse, the prevailing view is that dorsal horn CGRP derives exclusively from afferents. Here we took advantage of a tamoxifen-inducible CGRP^CreER^ mouse line, which when crossed with a tdTomato reporter mouse, reveals a discrete population of CGRP-expressing interneurons that are concentrated in lamina III and inner lamina II of the spinal cord dorsal horn and trigeminal nucleus caudalis.

Unlike dorsal horn vertical cells, which have ventrally-directed dendrites and a dorsally-directed axon, the CGRP interneurons have mainly dorsally-directed dendrites and ventrally-directed axons. A comprehensive functional analysis showed that these interneurons are minimally responsive to a host of acute, innocuous or noxious mechanical and chemical stimuli, despite the fact that electrical stimulation of Aβ afferents readily activates the cells. On the other hand, an innocuous mechanical stimulus evoked significant Fos expression in the setting of peripheral nerve injury and chemogenetic activation of the interneurons produced significant mechanical hypersensitivity. We conclude that these CGRP-expressing interneurons engage deep dorsal horn nociresponsive circuits that contribute to nerve injury-induced central sensitization and Aβ-mediated mechanical allodynia.

## RESULTS

To map the distribution of CGRP-expressing neurons in the dorsal horn, we first crossed the CGRP^CreER^ mouse line with a floxed stop ROSA-tdTomato line. Adult mice were administered tamoxifen twice (150 mg/kg, at postnatal days 21-23), and as reported previously, this triggered tdTomato expression in primary sensory neurons (Patil, Hovhannisyan, & Akopian, 2018). However, we also recorded significant labeling of neurons in the dorsal horn and trigeminal nucleus caudalis (N. Caudalis; Figure 1). Importantly, because the tamoxifen is administered at 3-4 weeks of age, we conclude that the pattern of expression is reflective of that found in the adult.

**Figure 1.**
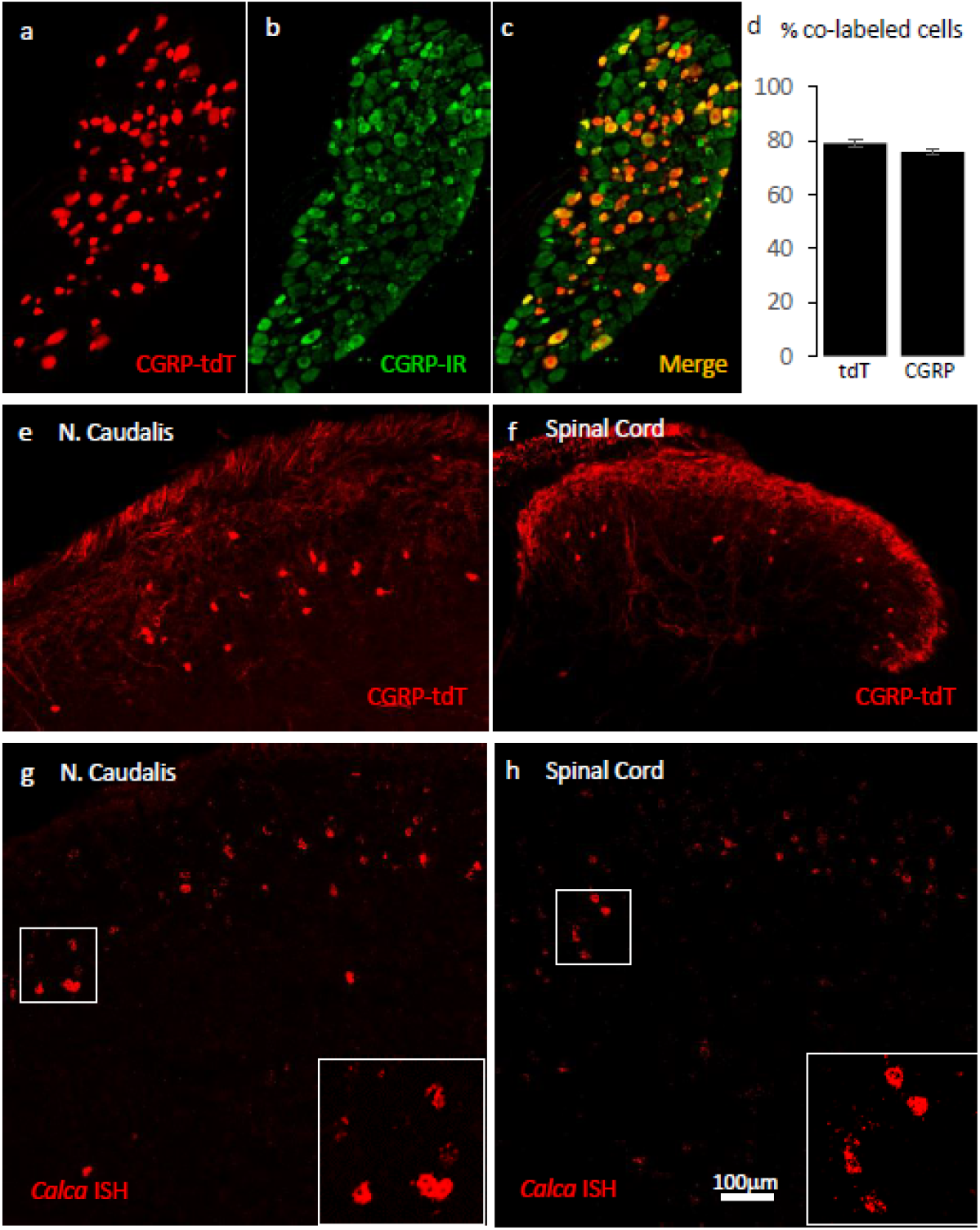
Validating the CGRP^Cre-ER^ transgenic mouse. **a-c)** Example of genetically labelled CGRP neurons from dorsal root ganglion of double transgenic CGRP^CreER^ / tdTomato mice generated by crossing the CGRP^CreER^ mouse line with a floxed stop ROSA-tdTomato line. Adult CGRP^CreER^ / tdTomato mice received 2 injections of tamoxifen (150 mg/kg). Co-localization of tdTomato-(red) with CGRP-immunoreactivity (green) confirmed the specificity of CGRP^CreER^ expression in trigeminal and dorsal root ganglia. **d)** 80% of tdTomato-positive neurons were immunoreactive for CGRP (left bar) and 78% of CGRP-positive neurons were tdTomato-immunoreactive (right bar). Bars show mean and standard error (SEM) (3 mice, 4 sections each). **e-f)** TdTomato expression was also detected in neurons of nucleus caudalis (**e)** and the spinal cord dorsal horn **(f)**. The tdTomato-immunoreactive neurons were concentrated in lamina III and occasionally observed in more superficial layers. **g-h)** *In situ* hybridization confirmed expression of CGRP mRNA in the dorsal horn **(g)** and nucleus caudalis **(h)**. Insets show higher magnification of the CGRP mRNA expressing neurons. Scale bars: 100 μm.

We first confirmed the approach by ensuring that the tdTomato-expressing primary sensory neurons of the dorsal root ganglia (DRG) double-label with an antibody to CGRP. Figure 1 illustrates that 80% of tdTomato-positive neurons in trigeminal ganglia (TG) and DRG immunostained for CGRP and that 78% of the CGRP immunoreactive neurons were tdTomato-positive (Figs. 1a-d).

Consistent with the central projection of CGRP–expressing sensory neurons, we also observed very dense tdTomato-positive terminals in the superficial laminae of the dorsal horn and nucleus caudalis. We also recorded many tdTomato-labeled neurons in regions of the central nervous system known to contain significant populations of CGRP-immunoreactive neurons or terminals, including motoneurons in the ventral horn of the spinal cord (Supplementary Figure 1), the parabrachial nucleus (Supplementary Figure 2), subparafascicularis of the thalamus (Supplementary Figure 3) (Yasui, Saper, & Cechetto, 1991), and central nucleus of the amygdala (Supplementary Figure 3) and in cranial motor nuclei (Supplementary Figure 4). We conclude that the pattern of CGRP-expression observed in the CGRP^CreER^ mouse provides a reliable marker of CGRP-expressing neurons in the adult.

**Figure 2.**
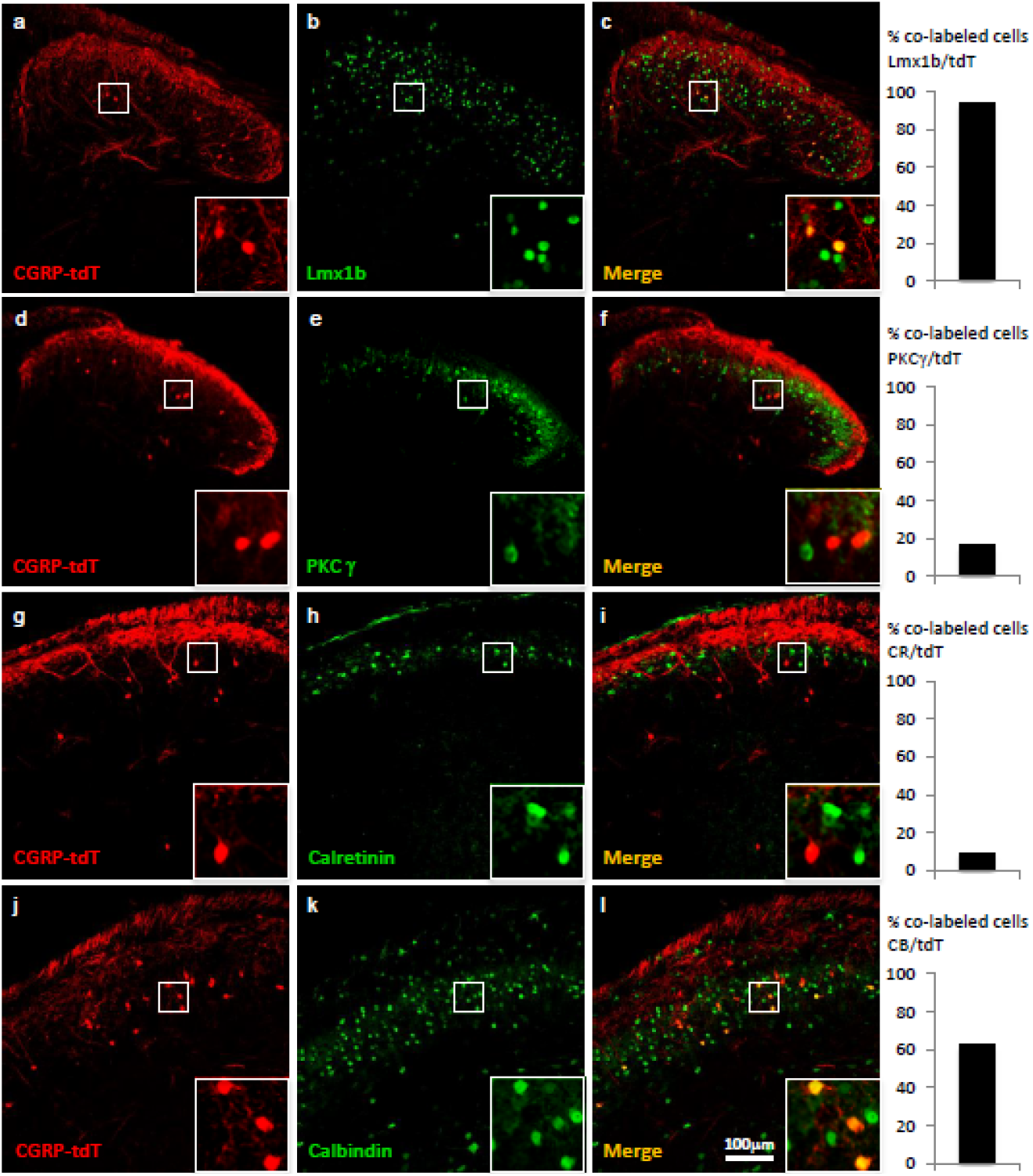
CGRP-expressing neurons in the dorsal horn (a-f) and nucleus caudalis (g-l) are a distinct class of excitatory (Lmx1b+) interneurons. **a-l)** Immunohistochemistry showed that CGRP-tdTomato fluorescent neurons (red) co-express many markers (green) of excitatory, but not inhibitory (e.g., Pax2, Supplementary Figure 5) interneurons in the dorsal horn **(a-f)** and nucleus caudalis **(g-l)**. Ninety-eight percent of CGRP-tdTomato neurons co-expressed Lmx1b (**a-c**), 16% co-expressed PKCγ (**d-f**), 9% co-expressed calretinin (**g-i**), and 63% co-expressed calbindin (**j-l**). Insets show higher magnification views of boxed areas in respective images. Graphs illustrate mean percentages ± SEM of CGRP-tdTomato neurons that were double-labelled with the indicated antibody (~100 cells per antibody). Scale bar: 100 μm.

**Figure 3.**
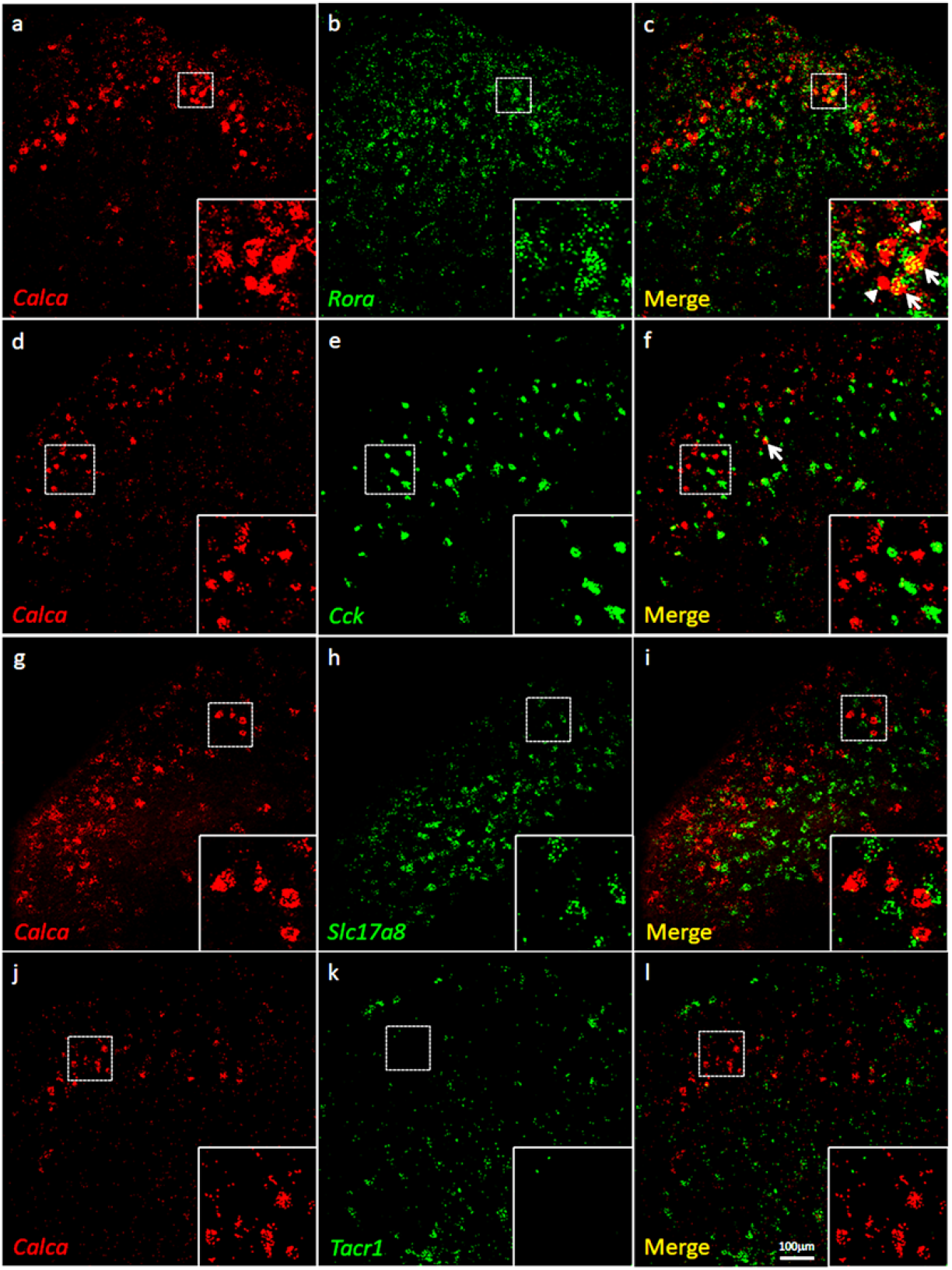
Coexpression of CGRP mRNA with RORα mRNA, but with neither CCK nor NK1R mRNA. **a-l)** Co-expression of CGRP mRNA (*Calca;* red), with other markers (green) in subsets of dorsal horn (**a-c; j-l**) and nucleus caudalis (**d-i**) neurons. Of *Calca*-expressing cells, 56% express RORα mRNA **(a-c)**, but only 4.4% express *Cck* mRNA **(d-f)**. Similarly, there was minimal overlap of *Calca* and *Slc17a8*, the gene coding for VGLUT3 **(g-i),** or *Calca* and *Tacr1*, the gene coding for the NK1 receptor **(j-l)**. Insets show higher magnification images of boxed areas. Scale bar: 100 μm.

**Figure 4.**
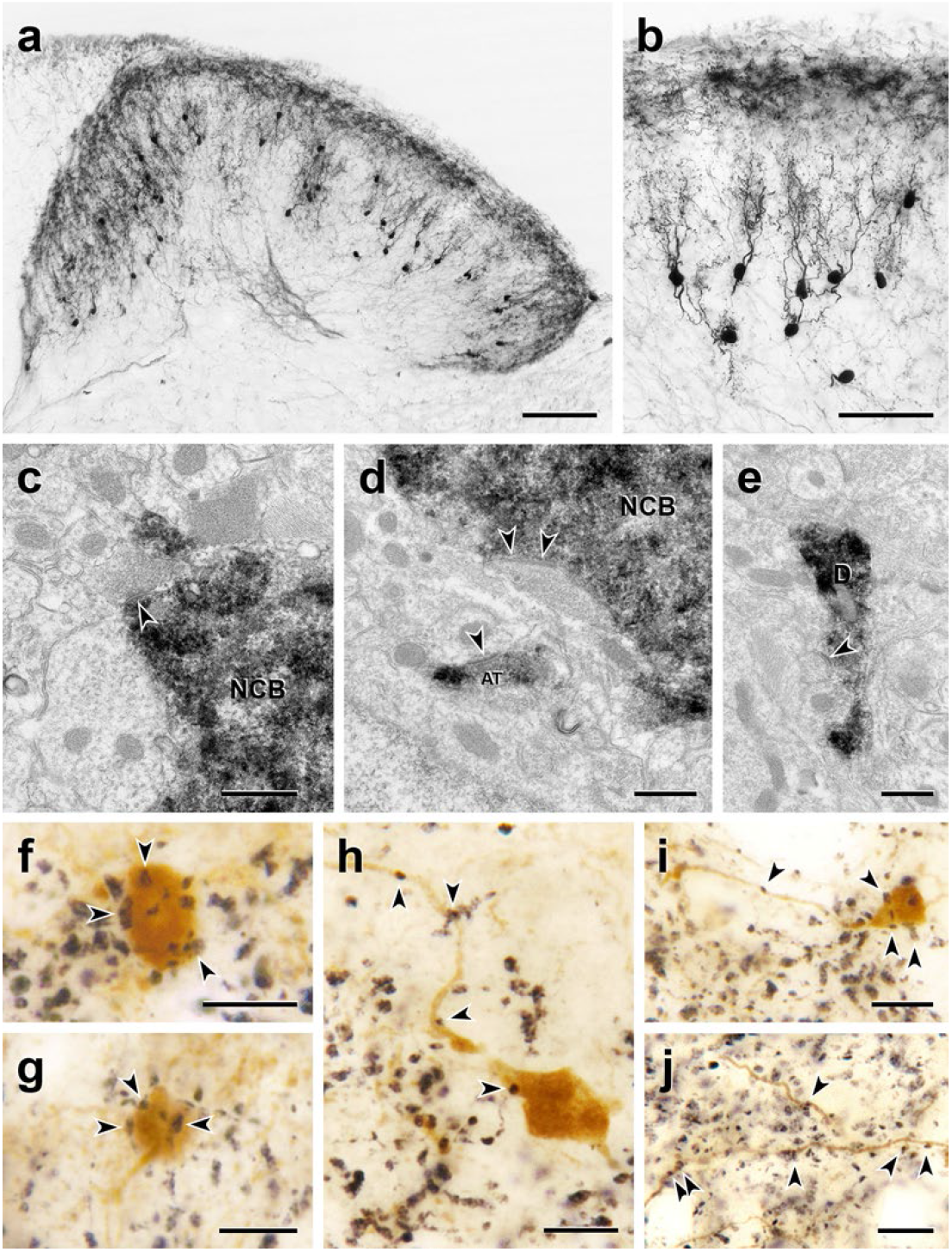
Morphology and VGLUT1 innervation of dorsal horn CGRP interneurons. **a, b**) Most tdT-immunoreactive CGRP interneurons (black) are located in lamina III and have a relatively uniform morphology with many spiny, dorsally-projecting dendrites. Scale bars: 100 μm in a; 50 μm in **b**. **c-e:** Electron microscopic analysis revealed unlabelled host synapses (arrowheads) presynaptic to the cell bodies (NCB in **c** and **d**) and dendrites (D in **e**) of tdT-immunoreactive (black) CGRP interneurons. **d** also shows an asymmetric presynaptic input (AT) from a presumptive CGRP interneuron to an unlabelled host dendrite. **f – j,** Black VGLUT1-immunoreactive varicosities form close appositions (arrowheads) with the cell bodies (**f & g**) and dendrites (**h – j**) of brown tdT-immunoreactive CGRP interneurons. Scale bars: 500 nm in **c** – **e**, 10 μm in **f** – **j**.

Unexpectedly, we also found large numbers of small tdTomato-positive neurons in the superficial dorsal horn and nucleus caudalis (notably in lamina III) and occasionally in more superficial layers (Figures 1e-f: Supplementary Figure 8c). Consistent with previous literature, we did not detect CGRP-immunoreactivity in dorsal horn neurons using well-validated antibodies. However, by *in situ* hybridization we confirmed that CGRP mRNA is present in neurons in the same regions of the spinal cord dorsal horn and nucleus caudalis (Figure 1g-h), which is consistent with the single cell PCR reports of CGRP message in subpopulations of dorsal horn neurons (Haring et al., 2018; Sathyamurthy et al., 2018). We speculate that the lack of CGRP immunostaining reflects rapid transport of the peptide from the cell body to its axon, which undoubtedly underlies the requirement for colchicine to demonstrate these neurons by immunocytochemistry (Kruger et al., 1988; Tie-Jun et al., 2001). We found the CGRP positive interneurons to be particularly abundant at the most caudal levels of the nucleus caudalis, markedly decreasing rostrally as the hypoglossal nucleus appears (Supplementary Figure 4).

### CGRP dorsal horn neurons are excitatory interneurons

We next asked whether these CGRP–expressing neurons include both projection and interneurons. First, we injected the retrograde tracer Fluorogold (1%) into several brain areas that receive projections from the spinal cord dorsal horn. Despite an extensive analysis, which included injections into the ventrobasal and nucleus submedius (Yoshida, Dostrovsky, Sessle, & Chiang, 1991) of the thalamus, lateral parabrachial nucleus, and dorsal column nuclei, which are targeted by postsynaptic dorsal column neurons located in the region of lamina IV of the dorsal horn, we found no evidence of CGRP-expressing projection neurons. This finding was confirmed with an anterograde tracing approach in which we injected an AAV1-flex-GCaMP6s virus unilaterally into the nucleus caudalis of CGRP^CreER^/tdT mice (Supplementary Figure 7). After 4 weeks, we examined the brainstem, thalamus and hypothalamus for GFP-labeled fibers, but found no evidence of long distance axonal projections deriving from the lamina III CGRP cells.

By immunolabeling the CGRP-tdTomato neurons, we next determined that these cells are excitatory and define a unique subset of interneurons. First, the CGRP-tdTomato cells co-express Lmx1b (98%; 92/94 tdT cells), but not Pax2 (Supplementary Figure 5), which are excitatory and inhibitory markers, respectively. Some of the CGRP-tdTomato cells populate inner lamina II, and here approximately 16% co-expressed PKCγ (31/187 tdT cells), a marker of a large population of excitatory interneurons (Malmberg, Chen, Tonegawa, & Basbaum, 1997). Sixty-three (97/158 tdT cells) and 9% (9/97 tdT cells) of the CGRP interneurons co-expressed calbindin and calretinin, respectively, calcium binding proteins that mark subpopulations of excitatory dorsal horn interneurons (Figure 2). The incomplete immunohistochemical overlap with major neurochemical classes of dorsal horn interneurons indicates that the CGRP interneurons are heterogeneous consistent with previously described populations of dorsal horn neurons. However, as there is a limited number of quality antibodies that can be used for comprehensive neurochemical profiling we turned to *in situ* hybridization (Figure 3).

**Figure 5.**
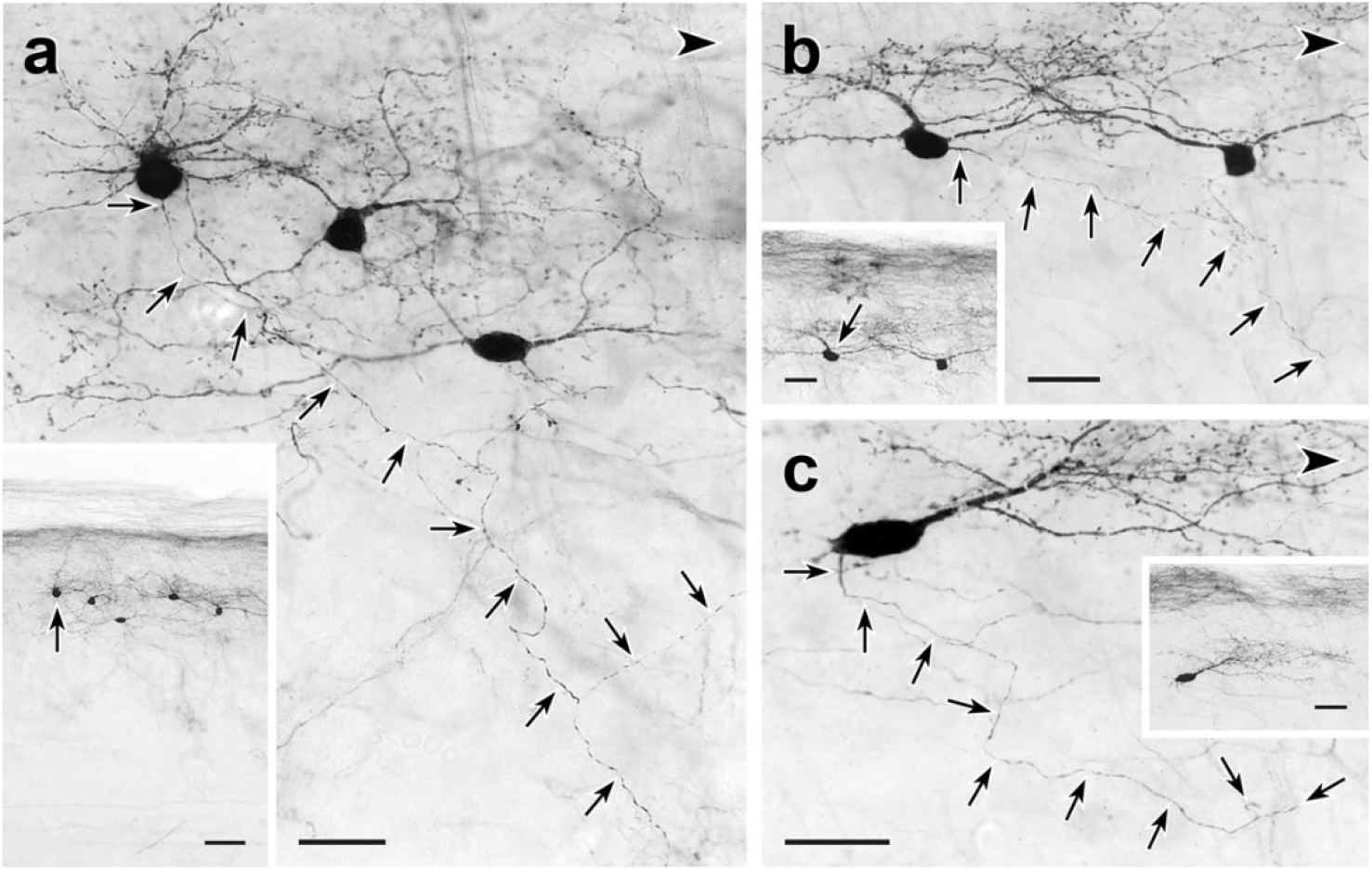
Trajectories of axons of CGRP-tdTomato interneurons. tdT-immunoreactive CGRP interneurons (black) in 50 μm parasagittal sections from the lumbar dorsal horn of CGRP-tdTomato mice in which an intrathecal injection of capsaicin reduced primary afferent-derived CGRP. The CGRP-tdTomato neurons have spiny, dorsally-directed dendrites and their axons (arrows) course ventrally and often caudally (large arrowhead). Arrows in insets indicate location of the neurons whose axons are shown in **a**, **b** and **c**. Scale bars: 20 μm in **a-c**, 50 μm in inset **a**, 20 μm in insets **b** and **c**.

Consistent with the concentration of tdTomato-CGRP interneurons in lamina III, particularly notable is that 56% of the CGRP mRNA-expressing (*calca*) cells double-labeled for RORα message (639/1134 CGRP mRNA-expressing cells), a marker of excitatory interneurons in lamina III (Bourane, Grossmann, et al., 2015). Interestingly, however, only 4% co-expressed CCK (27/595 CGRP mRNA-expressing cells), which marks a significant subset of the RORα population (Liu et al., 2018). As for other populations of excitatory interneurons, we found minimal overlap with the population that transiently expresses VGLUT3 (examined at P7) (Peirs et al., 2015) or others that express Nptx2, BDNF or the NK1 receptor, a marker of many projection neurons. We conclude that a substantial portion of the CGRP interneuron population overlaps with a subset of the CCK-negative RORα population of lamina III interneurons.

### CGRP interneurons have dorsally-directed dendritic arbors and are innervated by VGLUT1-expressing terminals

Despite the very intense tdTomato labeling of the cell bodies of the dorsal horn neurons, it was difficult to distinguish axonal processes from the dense primary sensory neuron-derived CGRP innervation. This was particularly the case when an antibody to tdTomato was used to detect the dorsal horn CGRP neurons. And unfortunately, although the cell body of the intracellularly recorded cells was readily filled with biotin dextran in electrophysiological slice preparations (see below), we never successfully filled dendrites or axons. Therefore, in a separate set of experiments we first reduced the complement of primary afferent-derived CGRP-derived by making an intrathecal injection of capsaicin, 7 days prior to perfusing the mice (Cavanaugh et al., 2009). In addition, tdTomato-immunoreactivity was revealed with immunoperoxidase staining so that sections could be analyzed by either light or electron microscopy (EM). The results from this approach were both striking and especially informative. Figure 4 illustrates that the CGRP interneurons have many dorsally-directed, spine-laden dendrites. These dendritic arbors often penetrated lamina II, and some labeled processes appeared to reach lamina I. Nevertheless, despite the capsaicin treatment, the latter were rare and difficult to distinguish from residual primary afferent-derived CGRP.

Based on their remarkably uniform dendritic morphology, the dorsal horn CGRP neurons appear to represent a subpopulation of excitatory, so-called radial interneurons (Grudt & Perl, 2002), however, the morphology of the CGRP-expressing radial interneurons differ considerably from those previously described in lamina II. First, the majority of lamina II radial cells have dendrites that arborize ventrally and axons that, if anything, project and collateralize dorsally, occasionally targeting presumptive projection neurons in lamina I. In contrast, not only do the CGRP interneurons have dorsally-directed dendrites, but almost all of their axons project ventrally and/or ventrocaudally. In some instances we could trace the axons well into the neck of the dorsal horn, including lamina V (Figure 5 and Supplementary Figure 6). Furthermore, EM analysis of these interneurons (Figure 5) illustrates that there is significant synaptic input to the soma, dendrites and spines of the CGRP interneurons. Finally, given the concentration of the CGRP interneurons in lamina III, we assumed that they receive primary afferent input from large myelinated afferents. Indeed when we double-immunostained for tdTomato and VGLUT1, a glutamate transporter that is highly expressed in large myelinated afferents (Oliveira et al., 2003), we observed many close appositions of VGLUT1-immunoreactive axon terminals onto the cell bodies and dendrites of the CGRP interneurons (Figure 4 f-j).

**Figure 6.**
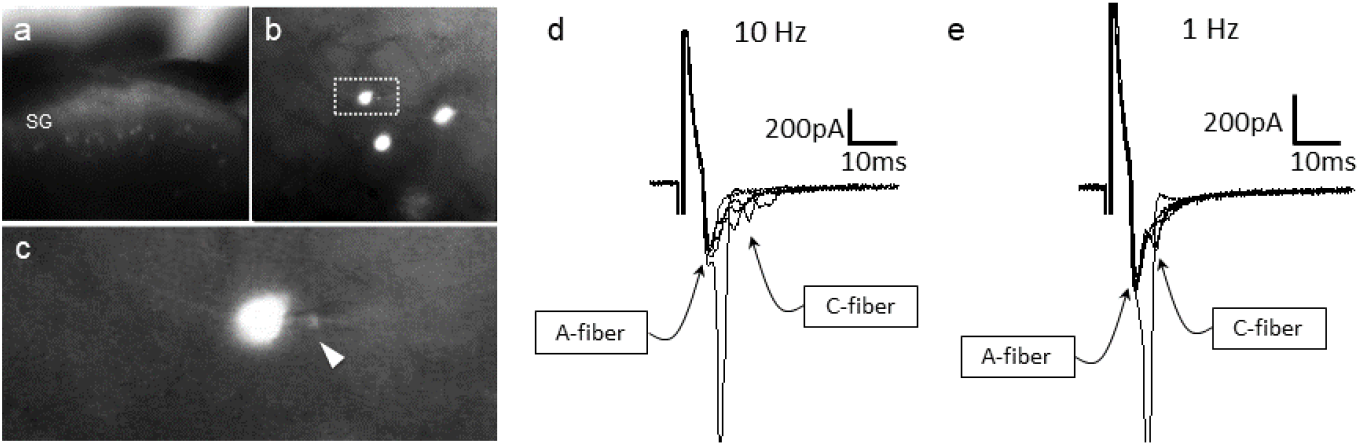
CGRP-tdTomato interneurons receive low threshold sensory inputs. Low (a) and high (b) magnification micrographs of endogenous fluorescent CGRP-tdTomato neurons in a spinal cord slice. The boxed neuron in **b** is shown at high magnification in **c** ; arrowhead points to the recording pipette in a whole cell configuration. **d,e)** Responses of CGRP-tdTomato interneuron to dorsal root stimulation at 10 Hz **(d)** or 1 Hz **(e).** An early, persistent component likely corresponds to a monosynaptic A-fiber input. The late component, with variable latency and failures, likely reflects polysynaptic C-fiber input.

**Figure 7.**
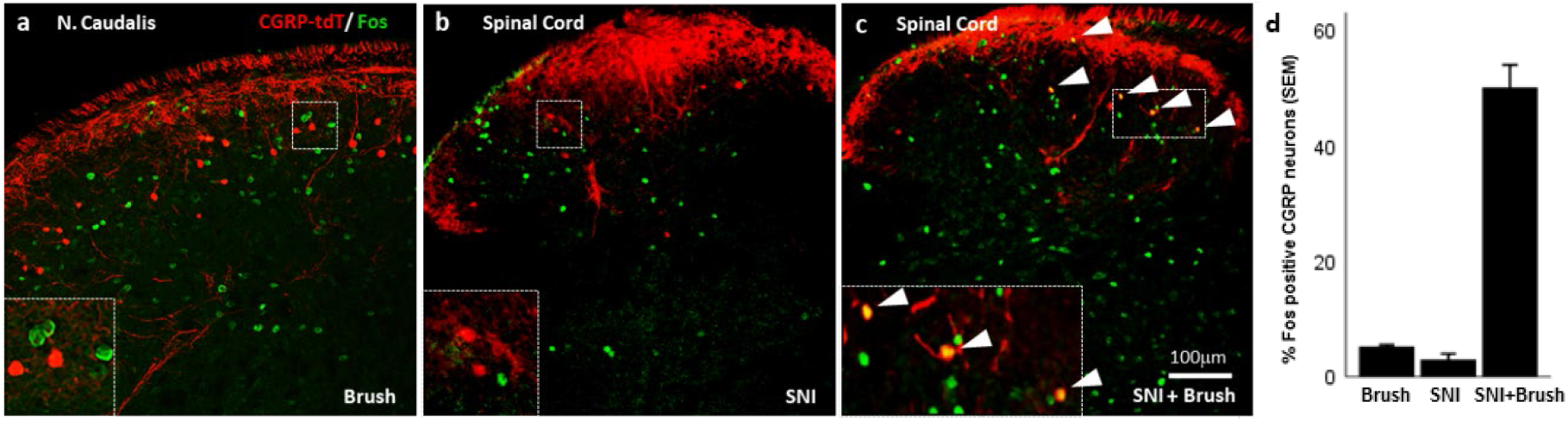
Peripheral innocuous stimuli activate CGRP interneurons but only after spared nerve injury (SNI). Fos-immunoreactive neurons in nucleus caudalis after brushing the cheek of a naïve uninjured mouse. **b)** Fos expression in the lumbar dorsal horn 6 day after SNI without additional peripheral stimulation. **c)** Fos expression in the lumbar dorsal horn 6-day after SNI with additional brush stimulation of the hindpaw. Insets: high magnification images of the boxed areas in the respective micrographs. Arrowheads indicate double-labeled cells. Scale bar: 100 μm. **d)** Mean percentages ± SEM of CGRP-tdTomato neurons that are Fos-immunoreactive in the different conditions.

**Figure 8.**
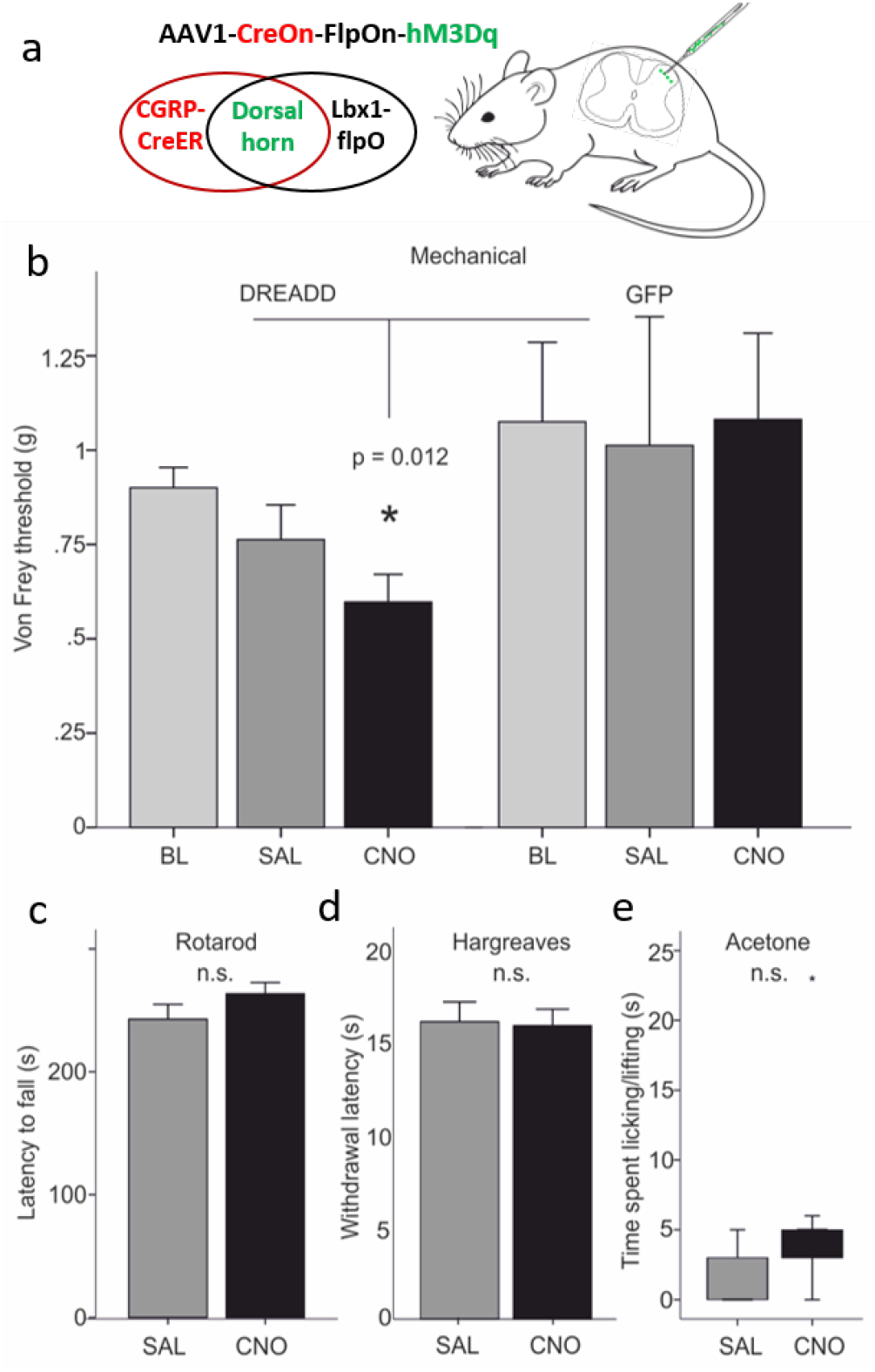
Dorsal horn CGRP interneurons contribute to mechanical hypersensitivity *in vivo*. **a**) CGRP^CreER^ mice were crossed to an Lbx1-driven FlpO mouse line, which restricts Cre expression to Lbx1-expressing neurons in the dorsal spinal cord and hindbrain. We then injected a Cre and Flp-dependent DREADD (hM3Dq) virus (AAV1-CreOn-FlpOn-hM3Dq) or a GFP-expressing AAV into the lumbar dorsal horn. **b)** Baseine (BL) von Frey mechanical thresholds of the DREADD-expressing mice (n=16; light grey bars) did not differ from baseline thresholds of mice injected with the AAV-GFP (GFP) control virus (n=6). In contrast, CNO injection significantly reduced von Frey thresholds (CNO, black bars) of the ipsilateral hindpaw in the DREADD-injected mice, compared either to their baseline, to the GFP controls or to saline (SAL; light grey bars)-injected mice (Repeated measures Two-way ANOVA, p=0.012). Neither latency to fall from a rotarod **(c)**, withdrawal to noxious heat in the Hargreaves test **(d)**, nor time spent paw lifting after exposure of the paw to a cold stimulus (acetone) **(e)** differed when comparing CNO and the control saline injection (p > 0.05, Students T-test and Wilcoxon Signed Ranks Test, respectively).

### CGRP-tdTomato interneurons receive low threshold primary afferent input

To confirm that the VGLUT1 appositions indeed mark a monosynaptic input from Aβ afferents to the CGRP-tdTomato interneurons, we prepared transverse lumbar and caudal medullary slices (350-400 μm) from 3-week old mice for whole-cell patch-clamp recordings. The slices contained large numbers of fluorescent tdTomato-labeled CGRP neurons (Figure 6a-c). We first characterized the intrinsic properties of the CGRP-tdTomato neurons by inducing depolarizing current steps. The CGRP-tdTomato neurons in the dorsal horn and nucleus caudalis showed mostly delayed firing patterns, consistent with their excitatory and radial phenotype (delayed 19, tonic 1, reluctant 2, single 2, no response 3, Supplementary Table 1). In some preparations we stimulated an attached dorsal root. At near threshold stimulation intensities (10 Hz), we recorded a very short latency component, which likely corresponds to a monosynaptic Aβ-fiber input. With more intense stimulation, we recorded a late component, with variable latency and failures. We assume that the latter derived from polysynaptic C-fiber input (Figure 6d-e). Of 5 cells recorded in 3 mice, all 5 received monosynaptic Aβ input and 2 of the 5 received polysynaptic C fiber input. In 2 additional mice we recorded from 4 cells that responded to dorsal root stimulation at A and/or C fiber threshold, but we could not unequivocally establish whether they received a monosynaptic input. Overall, the intrinsic properties of neurons recorded from lumbar dorsal horn (22 cells, 8 mice) and nucleus caudalis (5 cells, 2 mice) were comparable (see Supplementary Table 1). Taken together, we conclude that the predominant input to the CGRP interneurons derives from low threshold (Aβ) mechanoreceptors.

### Mechanical stimuli engage CGRP neurons, but only in a nerve injury setting

To provide a global activity measure of the stimuli that engage the CGRP interneurons, we first monitored Fos expression using a battery of noxious and innocuous stimuli. As expected, a unilateral injection of dilute formalin into the cheek (10 μl of 2% formalin, Supplementary Figure 8c) or a unilateral hindpaw injection of capsaicin (Supplementary Figure 9a-b), produced considerable Fos immunolabeling of dorsal horn neurons, but not of the CGRP-tdTomato interneurons (Supplementary Figures 8c and 9a-b). Unexpectedly, however, selectively engaging non-nociceptive afferents by having the animal walk for 90 minutes on a rotarod, which provokes considerable Fos in laminae III and IV (Neumann, Braz, Skinner, Llewellyn-Smith, & Basbaum, 2008), did not induce Fos expression in the CGRP interneurons (Supplementary Figure 8a). The same was true for brushing of the cheek, another innocuous stimulus that activates Aβ afferents (Figure 7). Finally, although CGRP is strongly implicated in the generation of migraine, largely but not exclusively via its peripheral vasodilatory action (Brain et al., 1985), systemic injection of nitroglycerin, which triggers migraine in humans and profound mechanical hypersensitivity in animals (Bates et al., 2010), did not induce Fos in the CGRP interneurons (Supplementary Figure 8b).

We conclude that despite our electrophysiological evidence that Aβ afferents engage the CGRP interneurons, there does not appear to be sufficient input to activate these cells under natural innocuous mechanical stimulus conditions in uninjured mice (5.3%, 5/88 tdT cells; Figure 7a). We, therefore, next asked whether an injury state would render the CGRP interneurons more responsive to an innocuous stimulus. In fact six days after inducing the spared nerve injury (SNI) model of neuropathic pain, we found that brushing the ipsilateral paw evoked Fos expression in 50% (53/110 tdT cells) of the dorsal horn CGRP interneurons (Figure 7c and d). Importantly, although we recorded significant dorsal horn Fos expression in nerve-injured mice without brushing (Figure 7b), no Fos expression occurred in the CGRP interneurons (3%; 6/205 tdT cells). We conclude that activation of the CGRP interneurons only occurs when the innocuous input engages the interneurons in the setting of nerve injury.

### Dorsal horn CGRP interneurons contribute to mechanical hypersensitivity *in vivo*

As electrical stimulation of the dorsal root at Aβ intensity readily excites the CGRP interneurons, the inability of brush stimulation to activate the neurons in the absence of injury was surprising. The discrepancy may reflect the fact that dorsal root stimulation involves a synchronous activation of many primary sensory neurons. In contrast, natural stimuli (e.g. brushing or walking on a rotarod) trigger an asynchronous afferent drive. However, as brushing was effective in the nerve injury setting, we hypothesized that a central sensitization rendered the CGRP neurons hyperexcitable. To test this hypothesis we asked whether a different synchronous stimulus, namely chemogenetic (direct) activation of the CGRP interneurons, could generate behaviors indicative of mechanical allodynia, comparable to what is observed in response to innocuous mechanical stimuli in the setting of nerve injury.

In these studies, we used an intersectional approach to target expression of a Designer Receptor Exclusively Activated by Designer Drugs (DREADD) selectively in the CGRP interneurons. To this end, we crossed the CGRP^creER^ mice to a FlpO mouse line, driven by the Lbx1 gene. The latter gene is only expressed in neurons of dorsal spinal cord and hindbrain, but not in sensory neurons of the DRG (Bourane, Grossmann, et al., 2015). We then made a unilateral microinjection of an adenoassociated virus (AAV) expressing a Cre and FlpO-dependent DREADD (hM3Dq) into the dorsal horn of the CGRP^CreER^/FlpO mice. Four weeks later we evaluated the behavioral effects of a systemic injection of CNO, which activates the DREADD.

We first established that there was no constitutive effect of virus infection. Thus, CNO injection, compared to saline, did not alter the latency to fall from an accelerating rotarod (Figure 8c). Furthermore, baseline von Frey mechanical thresholds of the DREADD-expressing mice, measured prior to injection of CNO, did not differ from mice injected with the AAV-GFP virus. In distinct contrast, Figure 8 shows that CNO injection in the experimental group produced a significant reduction of von Frey threshold of the ipsilateral hindpaw, compared to baseline or to saline-injected mice (Figure 8b). Mechanical thresholds did not change from baseline in the AAV-GFP control animals, whether they received saline or CNO (Repeated Measures Two-way ANOVA, F_(1,20)_=6.964, p=0.012, interaction effect between DREADD group and CNO treatment). The groups contained the same numbers of males and females (DREADD animals: 8 of each; GFP controls: 3 of each), but there was no significant interaction between sex and treatment (CNO versus saline). Nor did factoring in sex reduce the error (R^2^) in the full Repeated Measures Two-way ANOVA.

Lastly, we evaluated heat and cold responsiveness after CNO injection. Neither latency to withdraw the hindpaw to noxious heat in the Hargreaves test (*n* = 16; Figure 8d) nor time spent paw lifting after exposure of the plantar surface of the hindpaw to a cold (acetone) stimulus (*n* = 11; Figure 8e), differed when comparing CNO and control saline injection (p > 0.05, Students T-test and Wilcoxon Signed Ranks Test, respectively). We conclude that direct and synchronous activation of the CGRP interneurons produces a selective mechanical hypersensitivity, mimicking the mechanical allodynia observed in response to low threshold (Aβ) mechanical stimulation (brush) in the setting of nerve injury.

### CGRP interneurons and itch

Based on their single cell transcriptome analysis, Häring and colleagues (Haring et al., 2018) concluded that several populations of dorsal horn excitatory neurons that express CGRP mRNA co-express gastrin-releasing peptide (GRP), a peptide linked to dorsal horn circuits that drive itch-provoked scratching (Albisetti et al., 2019; Sun & Chen, 2007). To confirm this, we performed double *in situ* hybridization for CGRP and GRP. Although the GRP interneurons predominated in a band just dorsal to the CGRP interneurons, consistent with our previous report (Solorzano et al., 2015), we did find several instances of co-localization of CGRP mRNA and GRP mRNA. Interestingly, however, when using immunohistochemistry, we found almost no overlap of GRP and CGRP in a double transgenic GRP-GFP/CGRP-tdTomato mouse line (Supplementary Figure 10 a-d). Despite these discordant findings, we also examined the pattern of Fos expression provoked by injection of chloroquine (CQ), a strong pruritogen, into the cheek or hindpaw. To prevent scratching-induced Fos, the CQ injections were performed in anesthetized mice. As Supplementary Figure 10e-f illustrates, despite considerable chloroquine-induced Fos expression, we found only an occasional double-labeled neuron. We conclude that the CGRP interneurons, despite some overlap with GRP, likely do not transmit chemical itch, a finding consistent with the effects of deleting RORα (Bourane, Grossmann, et al., 2015). Whether the CGRP interneurons are engaged in conditions in which mechanical stimulation can trigger itch (alloknesis) remains to be determined.

## DISCUSSION

Despite overwhelming evidence that primary sensory neurons are the predominant source of dorsal horn CGRP, here we describe a morphologically uniform population of dorsal horn CGRP-expressing interneurons. Many of these interneurons correspond to the CCK-negative subset of the RORα population in lamina III of the dorsal horn and trigeminal nucleus caudalis, are excitatory and are activated by electrical stimulation of non-nociceptive, Aβ primary afferents. In contrast to the CCK-expressing subset of RORα neurons, and despite their location in the so-called, low threshold mechanorecipient zone of the dorsal horn (Abraira et al., 2017), the CGRP interneurons do not express Fos in response to natural Aβ-mediated, innocuous mechanical stimulation (brushing or walking on a rotarod). As for the RORα population, the CGRP interneurons do not respond to noxious chemical stimulation. Even peripheral nerve injury, without superimposed stimulation, did not activate these neurons. On the other hand, brush stimulation in the nerve injury setting did activate the CGRP interneurons. This distinction suggests that unless these neurons are rendered hyperexcitable, as occurs after nerve injury, only synchronous afferent input is sufficient to engage the circuits in which the CGRP interneurons participate. Consistent with this conclusion, chemically-provoked (chemogenetic) synchronous activation of these neurons produced a significant mechanical hypersensitivity. Based on the predominant ventrally-directed axonal arbors of these interneurons we suggest that the dorsal horn CGRP interneurons contribute either to ascending circuits originating in deep dorsal horn or to the reflex circuits through which nerve-injury induced mechanical allodynia is manifest.

RNA-Seq analyses have now defined at least 15 subsets of excitatory interneurons and 15 subsets of inhibitory neurons in the dorsal horn of the spinal cord (Haring et al., 2018; Sathyamurthy et al., 2018). Ablation, optogenetic and chemogenetic studies further characterized those classes based on functional properties. Of note, an increasing number of dorsal horn interneurons that “gate” mechanical pain have been identified. These include neurochemically distinct excitatory interneuron populations: transient VGLUT3, somatostatin, RORα, calretinin and Tac1 (Bourane, Duan, et al., 2015; Cheng et al., 2017; Duan et al., 2014; Huang et al., 2019; Peirs et al., 2015; Petitjean et al., 2019) and distinct inhibitory interneuron populations: dynorphin, calretinin, parvalbumin and encephalin (Boyle et al., 2019; Duan et al., 2014; Francois et al., 2017; Petitjean et al., 2019; Petitjean et al., 2015). The CGRP-expressing interneurons define yet another population of dorsal horn interneurons that contributes to spinal cord processing of mechanical inputs. Interestingly, there is a striking laminar organization of these molecularly distinct populations of interneurons. For example, the transiently expressing VGLUT3 population is located ventral to the CGRP interneurons, receives low-threshold mechanoreceptive input and their chemogenetic activation also enhances mechanical sensitivity (Cheng et al., 2017; Peirs et al., 2015). Dorsal to the CGRP interneuron are PKCγ and calretinin excitatory interneurons that contribute to nerve injury induced mechanical allodynia (Malmberg et al., 1997; Neumann et al., 2008; Peirs et al., 2015; Petitjean et al., 2019; Smith et al., 2019).

To what extent these mechanically-driven neuronal populations are interconnected or whether they represent parallel, independent circuits activated under different mechanical pain conditions (e.g. naïve vs injury vs inflammation) remains to be determined. Here the unique morphology of the CGRP interneurons is instructive. In contrast to many of the interneuron populations whose axons arborize longitudinally (e.g. PKCγ cells) or dorsally (e.g. calretinin cells), the CGRP interneurons have ventrally-directed axons. In some respects, the CGRP interneurons resemble the lamina II radial cells described by Grudt and Perl (Grudt & Perl, 2002) in the mouse, many of which are nociceptive, and the lamina III interneurons demonstrated in Golgi preparations in the cat and primate (Beal & Cooper, 1978; Maxwell, 1985). The fact that the CGRP interneurons show delayed firing patterns is also consistent with the properties of excitatory lamina II radial cells (Dickie et al., 2019; Grudt & Perl, 2002; Punnakkal, von Schoultz, Haenraets, Wildner, & Zeilhofer, 2014; Yasaka, Tiong, Hughes, Riddell, & Todd, 2010). Surprisingly, despite their dorsal dendrites, which extend into lamina II, where many nociceptive afferents terminate, we found no evidence that the CGRP interneurons are activated by acute noxious inputs (capsaicin or formalin). On the other hand, we did detect an occasional polysynaptic input following synchronous electrical stimulation of primary afferent C fibers. Most importantly, compared to the lamina II radial cells, we recorded much more extensive ventral axon trajectories of the CGRP interneurons, which suggests that these interneurons engage very different circuits in the dorsal and potentially ventral horn. In this regard, the CGRP interneurons are distinct from the calretinin interneurons that target lamina I projection neurons (Petitjean et al., 2019).

An RNA sequencing study of dorsal horn interneurons demonstrated expression of *calca*, the gene that encodes CGRP, in different clusters of neurons (Haring et al., 2018), including several that express *rora,* the gene that encodes RORα, Consistent with those results, our *in situ* hybridization studies found extensive co-expression of *calca* and *rora*. In fact, almost 55% of the CGRP interneurons co-express RORα message and there are significant similarities in their anatomical and functional properties (Bourane, Grossmann, et al., 2015). Specifically, the majority of RORα interneurons are excitatory and approximately 1/3 has a radial morphology, with ventrally arborizing axons. Furthermore, both the CGRP and RORα interneurons receive a monosynaptic Aβ afferent input and interestingly, despite the lack of response to capsaicin, some neurons in both populations receive a polysynaptic A delta and C input. Consistent with the report that deletion of the RORα population did not influence itch (Bourane, Grossmann, et al., 2015), and despite some overlap of the CGRP and GRP subsets of interneurons, we found that pruritogens did not activate the CGRP interneurons, namely did not induce Fos (Supplementary Figure 10). There are, however, some striking differences between the RORα and CGRP interneurons. For example, although a majority of the RORα interneurons co-express CCK, the CGRP interneurons rarely do. Furthermore, whereas RORα interneurons are activated by innocuous mechanical stimuli (e.g. brushing) in both naïve and injured conditions, the CGRP interneurons respond to innocuous stimuli only in the setting of nerve injury.

To our knowledge, the CGRP interneurons represent the first class of excitatory interneurons in lamina III that are unresponsive to innocuous mechanical stimulation under basal conditions despite receiving a monosynaptic Aβ input. One possibility is that the CGRP interneurons are tonically inhibited under normal conditions. Reduction of these inhibitory inputs in the setting of injury (Torsney & MacDermott, 2006) would render the neurons responsive to an innocuous stimulus (e.g. brush). In turn, the ventrally-directed axons of these interneurons could drive reflex withdrawal circuits, and/or engage ascending nociceptive pathways located in deep dorsal horn. The fact that DREADD-mediated direct activation of many CGRP interneurons lowered mechanical withdrawal thresholds is consistent with that hypothesis. In other words, we suggest that sensitization of these neurons is critical to mechanisms that underlie Aβ-mediated mechanical allodynia in the setting of nerve injury. Interestingly, Lu et al (Lu et al., 2013) provided evidence for convergence of a primary afferent-derived Aβ and a tonic glycinergic inhibitory input to PKCγ interneurons, some of which we found express CGRP. Loss of this glycinergic inhibition allowed Aβ input to access lamina I nociceptive circuits. Other studies demonstrated a comparable outcome, in this case by a presynaptic glycinergic inhibition of non-nociceptive inputs to superficial dorsal horn neurons (Sherman & Loomis, 1996). Furthermore, Imlach et al (Imlach, Bhola, Mohammadi, & Christie, 2016) proposed that decreased glycinergic inhibition is selective for radial cells in lamina II and likely contributes to neuropathic pain. We suggest that a comparable circuit involving the CGRP radial cells could uncover low threshold inputs to ventrally located nociceptive circuits, which in recent years have been largely ignored (Wercberger and Basbaum, 2019).

## MATERIALS AND METHODS

### Animals

Mice were housed in cages on a standard 12:12 hour light/dark cycle with food and water ad libitum. Permission for all animal experiments was obtained and overseen by the Institutional Animal Care and Use Committee (IACUC) at the University of California San Francisco. All experiments were carried out in accordance with the National Institutes of Health Guide for the Care and Use of Laboratory Animals and the recommendations of the International Association for the Study of Pain.

### Mouse strains

The CGRP^CreER^ mouse strain was kindly provided by Dr. Pao-Tien Chuang (UC San Francisco) (Song et al., 2012). CGRP^CreER^ mice were then bred with C57BL/6J -Ai14 mice (Jackson Laboratory, Stock No: 007914) or with mice that selectively express green fluorescent protein (GFP) in gastrin-releasing peptide (GRP)-expressing cells (GRP-GFP mouse (Solorzano et al., 2015)). Lbx1FlpO mice, in which FlpO is driven from the Lbx1 promoter, were a kind gift from Dr. Martin Goulding at the Salk Institute, La Jolla CA.

### Tamoxifen

We dissolved tamoxifen (T5648, Sigma-Aldrich) in corn oil and injected it (150 mg/kg, i.p.) into the CGRP-tdTomato mice on two consecutive days. For immunohistochemistry, electrophysiology and tracing experiments we injected the tamoxifen into P21-22 mice. We waited 5 and 7-10 days before recording and perfusion for immunostaining, respectively. For Fluorogold (1%) tracing experiments, we injected the tracer into 6-8 week old mice. For intraspinal surgeries intended for DREADD receptor expression studies, we injected tamoxifen into P11-12 mice and subsequently, between P14-16, made an intraspinal injection of hM3Dq without laminectomy.

### Fluorescence Immunohistochemistry (IHC)

Mice of either sex were transcardially perfused with 10 mL phosphate-buffered saline (PBS) followed by 30 mL cold 4% formaldehyde in PBS. After dissection, dorsal root ganglia (DRG), trigeminal ganglia (TG), spinal cord and caudal medullary tissue were post fixed for ~3 hours at room temperature and subsequently cryoprotected in 30% sucrose in PBS overnight at 4°C. The spinal cord and caudal medulla were sectioned in a cryostat at 25 μm; DRG and TG at 16 μm. After mounting and drying on slides, the sections were incubated for 1.5 hours in 10% normal goat serum with 0.3% Triton X-100 (NGST) to block non-specific antibody binding, and then for 24 hours in primary antibodies diluted in 10% NGST. The sections were then washed 3 times for 10 minutes in PBS and then incubated for 2 hours with a secondary antibody diluted in 1% NGST. After washing with PBS 3 times for 10 minutes, the sections were dried and coverslipped with Fluoromount G.

The following primary antibodies were used: rabbit anti-CGRP (1:1000, Peninsula), rabbit anti-calbindin (1:2000, Swant), mouse anti-calretinin (1:5000, Swant), guinea pig anti-PKCγ (1:7000, Strategic Bio), chicken anti-GFP (1:2500, Abcam), rabbit anti-Fos (1:5000, Calbiochem; 1:2000, Cell Signaling), guinea pig anti-Fluorogold (1:1000, Protos Biotech), guinea pig anti-Lmx1b (1:10000, kind gift from T. Müller and C. Birchmeier, Max Delbrück Center for Molecular Medicine, Berlin, Germany), rabbit anti-Pax2 (1:4000, Abcam), or rabbit anti HA (1:800, Cell Signaling). Secondary antibodies were conjugated to Alexa-488 or Alexa-647 (1:1000, Thermo Fisher Scientific).

### Capsaicin treatment

To ablate the central terminals of CGRP-expressing DRG neurons, CGRP-tdTomato mice were anesthetized with 2% isoflurane and injected intrathecally with 5.0 μl of a solution containing 10 μg of capsaicin, dissolved in 10% ethanol, 10% Tween-80 and 80% saline. Five days later, the mice received 5 i.p. injections of 150 mg/kg tamoxifen (one injection per day, on 5 consecutive days). Seven days later, the mice were processed for immunohistochemistry.

### Peroxidase immunocytochemistry

Mice were perfused with phosphate-buffered 4% formaldehyde (n=3) or 4% formaldehyde plus 0.3% glutaraldehyde (n=5). Transverse or parasagittal Vibratome sections (50 μm) were processed for detection of tdTomato for either light (LM) or electron microscopic (EM) (Llewellyn-Smith, Basbaum, & Braz, 2018) examination.

### Electron microscopy

For EM analysis, the sections were washed for 2 h in 50% ethanol, incubated for 30 min in 10% normal horse serum diluted with Tris-PBS (TPBS), then in 1:25,000 or 1:100,000 rabbit anti DSRed (Takara Bio USA) in 10% NHS-TPBS. The sections were subsequently exposed to 1:500 biotinylated donkey anti-rabbit IgG (Jackson ImmunoResearch) in 1% NHS-TPBS and then to 1:1500 ExtrAvidin-horseradish peroxidase (Sigma-Aldrich) in TPBS. Incubations in immunoreagents were for 3 days at room temperature on a shaker; sections were washed 3 × 30 min between incubations. To visualize CGRP-tdTomato-immunoreactivity in the dorsal horn, we used a nickel-intensified diaminobenzidine (DAB) reaction and hydrogen peroxide generated by glucose oxidase (Llewellyn-Smith, Dicarlo, Collins, & Keast, 2005). After the peroxidase reaction, sections containing tdTomato-immunoreactive neurons were osmicated, stained *en bloc* with aqueous uranyl acetate, dehydrated with acetone and propylene oxide, and infiltrated with Durcupan resin (Sigma-Aldrich). Finally, sections were embedded on glass slides under Aclar coverslips (Electron Microscopy Sciences) and polymerized at 60°C for at least 48 hr. Dorsal horn regions containing CGRP-tdTomato neurons were re-embedded in resin on blank blocks under glass coverslips and repolymerized. Ultrathin sections were collected on copper mesh grids, stained with aqueous uranyl acetate, and examined with a JEOL 100CXII transmission electron microscope.

### LM analysis

Transverse or parasagittal Vibratome sections of tissue from mice perfused with phosphate-buffered 4% formaldehyde (n=3) or 4% formaldehyde, 0.3% glutaraldehyde (n=3) were either single stained to show tdTomato-immunoreactivity or double stained to demonstrate the relationships between VGLUT1-immunoreactive axons and CGRP-tdTomato neurons. All sections were washed 3 x 20 min in TPBS containing 0.3% Triton X-100 and exposed to 10% NHS in TPBS-Triton for 30 min. Single labeling involved exposure of sections to 1:25,000 or 1:100,000 anti-DSRed (Takara), 1:500 anti-rabbit IgG, 1:1500 ExtrAvidin-HRP and a nickel-intensified DAB reaction. For double labeling, VGLUT1-immunoeractivity was first detected with 1:50,000 or 1:100,000 rabbit anti-VGLUT1 (Synaptic Systems), biotinylated donkey anti-rabbit IgG, ExtrAvidin-horseradish peroxidase and a cobalt+nickel-intensified DAB reaction (Llewellyn-Smith et al., 2005). Then, after another blocking step in 10% NHS, DSRed-immunoreactivity was detected as for single labeling except that the peroxidase reaction was intensified with imidazole (Llewellyn-Smith et al., 2005) rather than nickel. For LM labeling, primary antibodies were diluted with 10% NHS in TPBS-Triton; secondary antibodies, in 1% NHS-TPBS-Triton; and avidin-HRP complex, in TPBS-Triton. For LM, all incubations in immunoreagents were done on a shaker at room temperature for at least 24 hours and washes between incubations were 3 x 20 min in TPBS. Stained sections were mounted on subbed slides, dehydrated and coverslipped with Permaslip Mounting Medium (Alban Scientific).

### *In situ* hybridization (ISH)

*In situ* hybridization was performed using fresh spinal cord or caudal medullary tissue from adult mice (8–10-week-old), except for transient VGLUT3 assessment (Peirs et al., 2015), where the mice were 7 days old. We followed the protocol outlined by Advanced Cell Diagnostics (Newark, CA). The tissue was dissected out, instantaneously frozen on dry ice, and kept at –80°C until use. Cryostat sections of DRG (12 μm) were fixed at 4°C in 4% formaldehyde for 15 min, washed twice in PBS, and dehydrated through successive 5 minute ethanol steps (50%, 70%, and 100%) and then dried at room temperature. After a 30 min incubation with protease IV, sections were washed twice in PBS and incubated at 40°C with RNAscope-labeled mouse probes: calcitonin gene-related peptide (CGRP), RAR-related orphan receptor alpha (RORα), cholecystokinin (CCK), vesicular glutamate transporter 3 (VGLUT3), neurokinin receptor 1 (NK1R), gastrin releasing peptide (GRP) for 2 h in a humidified chamber. Sections were then washed twice in washing buffer and incubated with four 15-30 minute “signal amplifying” solutions at 40°C. After two washes, the sections were dried and covered with mounting media containing 4′,6-diamidino-2-phenylindole (DAPI).

### Image analysis

Images of fluorescent immunostained sections were acquired on an LSM 700 confocal microscope using ZEN Software (Carl Zeiss). The microscope was equipped with 405, 488, 555, and 639 nm diode lasers. For co-localization studies we used a 20× Plan-Apochromat (20×/0.8) objective (Zeiss) and image dimensions of 1024 × 1024 pixels with an image depth of 12 bits. Two times averaging was applied during image acquisition. Laser power and gain were adjusted to avoid saturation of single pixels and kept constant for each experiment. Image acquisition was performed with fixed exposure times for each channel and a 10% overlap of neighboring images where tiling was used. Stitching was done in ZEN using the “stitching/fuse tiles” function. Adjustment of brightness/contrast and maximum projections of Z-stack images were done in Fiji/Image J. All images of the same experiment were processed in an identical manner.

Images of peroxidase immunostained sections were acquired on an Olympus BH2 brightfield microscope equipped with SPlanApo lenses and a SPOT Insight CMOS Color Mosaic 5MP camera running SPOT 5.3 Advanced software. For assessment of VGLUT1 appositions on DSRed-immunoreactive CGRP neurons, an ×100 oil immersion lens was used. A VGLUT1-positive terminal was classified as forming a close apposition when (1) there was no space between the terminal and the DSRed-positive neuron for terminals lying side-by-side with a cell body or dendrite or when (2) the terminal and the DSRed-positive neuron were in the same focal plane for terminals overlying cell bodies or dendrites.

### Cell counts

To analyze overlap by immunohistochemistry or *in situ* hybridization, we counted cells from 4-5 sections in at least 3 animals per experiment. By immunohistochemistry, we first counted the number of neurons in the DRG and TG that were tdTomato-positive (total 1266 cells, 3 mice) or CGRP-positive (total 1050 cells, 3 mice) and then determined the percentage of tdTomato-positive neurons that were CGRP double-labeled and vice versa. The number of dorsal horn tdTomato-positive cells that double-labeled for different markers (e.g. PKCγ, Lmx1b, Fos, calretinin, calbindin) are indicated in the Results. To conclude that cells were double-labeled by *in situ* hybridization we set a threshold of at least 5 fluorescent “dots” for each probe in conjunction with a DAPI-positive nucleus.

### Viral vectors

For DREADD experiments we used a Cre and FlpO-dependent hM3D(Gq) adeno-associated virus: AAV1--hEF1alpha/hTLV1-Fon/Con[dFRT-HA_hM3D(Gq)-dlox-hM3D(Gq)-I-dlox-I-HA_hM3D(Gq)(rev)-dFRT]-WPRE-hGHp custom made by the University of Zurich Viral Vector Facility of the Neuroscience Center. For control injections we used an AAV1.hSyn.eGFP.WPRE.SV40 from Addgene. For GCaMP-tracing experiments we used an AV1.Syn.Flex.GCaMP6s.WPRE.SV40 from the Penn Vector Core, University of Pennsylvania. Note that we evaluated several Cre-dependent viral vectors for the tracing studies and only used those where specificity of expression was confirmed by lack of expression after injection into WT mice. We waited at least 4 weeks to achieve stable viral expression before beginning the behavioral or neuroanatomical experiments.

### Retrograde tracing

To study potential projection targets of the dorsal horn CGRP interneurons, we injected Fluorogold (1%) into several supraspinal sites known to receive projections from the spinal and medullary dorsal horns. We studied two mice for each location and allowed 5-9 days for tracer transport after which the mice were perfused with formaldehyde for subsequent histological analysis. We injected tracer into the following locations: ventrolateral thalamus (X:ML=1.5, Y:AP=−1.82, Z:DV=3.5; 500 or 800 nl); parabrachial nucleus (X=1.25, Y=−4.95, Z=−3.6; 600 nl); nucleus submedius of the thalamus (X=0.5, Y=−1.43, Z=4.25; 250 or 450 nl): dorsal column nuclei (400 nl).

### AAV injections

For all surgeries, the mice were administered carprofen (0.1 mg/kg, i.p.) just prior to surgery and lidocaine (0.5%) was applied to the incision site. For the DREADD experiments, under 2% isoflurane anesthesia we injected P14-16 CGRP^creER^-LbxflpO mice and littermates with an AAV-GFP. We removed muscles that overlay the left side of the T13 and L1 vertebra to expose the lumbar enlargement. Without laminectomy, we then slowly inserted a glass micropipette (50 μm tip) through the dura and made two 400 nl rostrocaudally separated injections of viral solution. The micropipette was left in place for ~2 minutes after which overlying muscle and skin were closed. After recovering from the anesthesia, the mice were returned to their home cages.

For the GCaMP6 tracing studies we made injections (300 – 800 nl) into the medullary dorsal horn in 8 week old mice anesthetized with i.p. ketamine (100 mg/kg) and xylazine (10 mg/kg) or isofluorane (2%). For injections into the nucleus caudalis, we incised the dura overlying the cisterna magna, exposing the caudal medulla and made a unilateral injection of viral solution with a glass micropipette. After recovering from anesthesia the mice were returned to their home cage.

### Behavioral analyses

We took several measures to blind the behavioral experiments. 1) DREADD-injected and control (GFP-injected) mice were housed together. 2) A different experimenter performed the injections of CNO (5.0 mg/kg in saline) or saline before behavioral testing. 3) The behavioral tester recorded each mouse’s eartag number after the test and was blind to the treatment (saline or CNO) that the mouse received or to which group the mouse belonged (AAV-GFP-injected or DREADD-injected). 4) Identification was made using records of eartag numbers after all testing was finalized.

#### Static mechanical allodynia

For these experiments, we determined hindpaw mechanical thresholds with von Frey filaments, and quantified results using the updown method (Chaplan, Bach, Pogrel, Chung, & Yaksh, 1994). The animals were habituated on a wire mesh for 2 hours on 2 consecutive days. On the next 2 days we recorded baseline thresholds, after a 1.5 hours of acclimatization on the wire mesh. After baseline determinations, the mice were injected with CNO or saline and then tested 30 minutes later. For all behavioral tests, either CNO or saline was injected every other day in a randomized fashion.

#### Acetone test (cold allodynia)

Mice were habituated for 30 minutes on a mesh in plexiglass cylinders. Next we used a syringe to squirt 50 μl acetone onto the plantar surface of the paw. The responses of the mice directly after application of acetone were recorded on video for 30 seconds. Each paw was tested 5 times and we measured time (in seconds) spent lifting, licking or flinching the paw. Results are displayed as the average time across the 5 trials. Testing began 1 hour post injection of CNO or saline, with test days 48 hours apart.

#### Hargreaves test

For thermal threshold testing (heat), we first acclimatized the mice for 30 minutes in Plexiglass cylinders. The mice were then placed on the glass of a Hargreaves apparatus and the latency to withdraw the paw from the heat source was recorded. Each paw was tested 5 times and we averaged latencies over the 5 trials. Hargreaves tests were done 1 hour after the tests of static dynamic mechanical allodynia.

#### Rotarod test

Mice were acclimatized to the testing room and trained by placing them on an accelerating rotarod for a maximum of 60 sec at low speed, 3 times with training taking place on two consecutive days. On testing days (48 hours apart), mice were injected with CNO or saline 30 minutes before being placed on the rotarod. Latency to fall was measured for up to 300 seconds. The procedure was repeated 3 times and latencies averaged across trials.

### Spared nerve injury (SNI)

To induce mechanical hypersensitivity in a model of neuropathic pain we used the approach described by Shields et al. (Shields, Eckert, & Basbaum, 2003). Under isofluorane anesthesia (2%), two of the three branches of the sciatic nerve were ligated and transected distally, sparing the tibial nerve.

#### Fos expression: Capsaicin and formalin

To study the effects of a chemical algogen, we injected 10 μl of 2% formalin in saline into the cheek (n=3). In a separate group of anesthetized animals (n=3), we made a unilateral injection of 20 μl capsaicin (1.0 μg/μl) into the hindpaw or the cheek. We perfused all mice ~1.5 hours after injection and immunostained sections of the lumbar cord (paw injections) or caudal medulla (cheek injections) for Fos.

#### Fos expression: Chloroquine

To study the effects of a pruritogen, under isofluorane anesthesia, mice (n=3) received unilateral injections of chloroquine (200 μg) into either the hindpaw (20 μl) or cheek (50 μl). The mice were perfused ~1.5 hours after injection and sections of the lumbar cord (paw injections) or caudal medulla (cheek injections) were immunostained for Fos.

#### Fos expression: Nitroglycerin

We injected mice (n=3) with nitroglycerin (10 mg/kg, i.p.), which in humans can trigger a migraine and in rodents provokes behavioral signs of widespread thermal hyperalgesia and mechanical hypersensitivity (Bates et al., 2010), beginning 30-60 min after injection and subsiding within 4 hours. Based on this time course, the mice were perfused 2 hours after nitroglycerin injection and sections caudal medulla were immunostained for Fos.

#### Fos expression: Dynamic mechanical allodynia

To assess Fos expression in uninjured animals (n=3), we first acclimatized the mice to brushing of the cheek, (Utrecht 225, pure red sable brush 6, Germany) while lightly restraining the mouse in a towel with its head exposed. We brushed the left cheek along the direction of the hairs for 45 min, with a one minute break every 10 minutes. To monitor Fos expression in the injured animals, we performed unilateral partial sciatic nerve injury (SNI, see above). One week after SNI, we used a paintbrush (5/0, Princeton Art & Brush Co.) to lightly stroke the injured hind paw, from heel to toe (velocity: ~2 cm/s). Ninety minutes to 2 hours after brushing, the mice were anesthetized, perfused and spinal cord sections were immunostained for Fos. In a separate experiment, we also assessed Fos expression one week after SNI without applying a stimulus.

#### Fos expression: Rotarod test

Three mice were trained on a rotating rod for 60 min at a constant speed of 10 rpm. One week later the mice walked on the rotarod at 10 rpm for 1.5 hours (Neumann et al., 2008), after which they were anesthetized, perfused and lumbar spinal cord sections immunostained for Fos.

### Electrophysiology

Following our previous protocol (Etlin et al., 2016), we collected transverse lumbar and caudal medullary Vibratome (Leica) slices (350-400 μm) from 3-10 weeks old CGRP-tdT mice 5-7 days after tamoxifen injection. The sections were incubated in recording solution at 37°C for 1 hour and then transferred to a recording chamber (Automate Scientific) under an upright fluorescence microscope (E600FN; Nikon). The sections were superfused with recording solution at a rate of 1.0 ml/min and viewed with a CCD digital camera (Hamamatsu or DAGE-MTI). The transparent appearance of lamina II of the superficial dorsal horn and tdTomato-positive CGRP cells were obvious under near-infrared (IR) illumination. The patch pipettes were pulled to yield an impedance of 6–8 MΩ on a horizontal pipette puller (Sutter Instrument) from thin-walled, fire-polished, borosilicate glass filaments. The pipette solution composition was (in mM): K-methane sulfonate 140, NaCl 10, CaCl_2_ 1.0, EGTA 1.0, HEPES 10, Mg-ATP 5.0, and NaGTP 0.5 and included 5.0 mg/ml of Biocytin (Sigma-Aldrich) for intracellular filling of the recorded cells. Neurons were approached with a micromanipulator (Sutter Instrument) while monitoring the resistance in voltage-clamp mode using the “Membrane Test” module of pClamp10 software (Molecular Devices). To prevent clogging of the tip, we applied positive pressure to the pipette via a 1.0 ml syringe. After a seal was established with a cell, we ruptured its membrane by gently applying negative pressure to the pipette to secure a whole-cell configuration. Current and voltage signals were amplified using a DC amplifier (MultiClamp 700) and digitized using Digidata 1440a system (Molecular Devices) at 10 kHz and then stored for subsequent offline analysis. In some experiments, we placed an attached dorsal root in a suction electrode to be stimulated electrically while simultaneously measuring evoked responses of the tdTomato-expressing neurons.

### Statistical analysis

Statistical analyses were performed using SPSS (IBM-SPSS version 24). Similarity of normality and variance were assessed before applying parametric or non-parametric tests. For analysis of the effect of CNO on mechanical hypersensitivity, we assessed interaction between treatment (CNO, saline or baseline) with group (DREADD-virus injected animals or GFP-virus injected animals) by repeated measures two-way ANOVA, including all conditions and groups. Statistics were calculated based on a type III sum of squares model and significant interaction effects were assessed using deviation from the mean of the control groups. The N was estimated based on variance for von Frey experiments using an *a priori* power calculation. Hargreaves and rotarod results were analyzed using Student’s t-tests. For acetone sensitivity we used the Wilcoxon signed rank test. Parametric and non-parametric tests are reported as mean +/− SEM or by medians and inter-quartiles, respectively. Electrophysiological recordings of intrinsic membrane and action potential properties were calculated using custom-written Matlab scripts (MathWorks, Illinois) as previously described (Etlin et al., 2016). P values were considered significant if p < 0.05.

## Acknowledgements

This research was supported by a Sir Henry Wellcome Fellowship 092208/Z/10/Z (LSL), NIH: R35NS097306 (AIB) and Wellcome Award: A102645 (AIB). We are grateful to Dr. Hendrik Wildner, University of Zurich for sharing the Cre/Flp dependent DREADD construct and to Dr. Ling Bai, University of California San Francisco for helpful advice on surgeries.

## Author contributions

LL, JMB and AIB conceptualized and designed the study. LL, AE, MB, MS, JK, KH, IL-S and JMB performed the experiments and collected the data. LL, AE, IL-S, JMB and AIB analyzed data. LL, JMB, AE, IL-S and AIB wrote the manuscript.

## SUPPLEMENTARY FIGURES

**Supplementary Figure 1.**
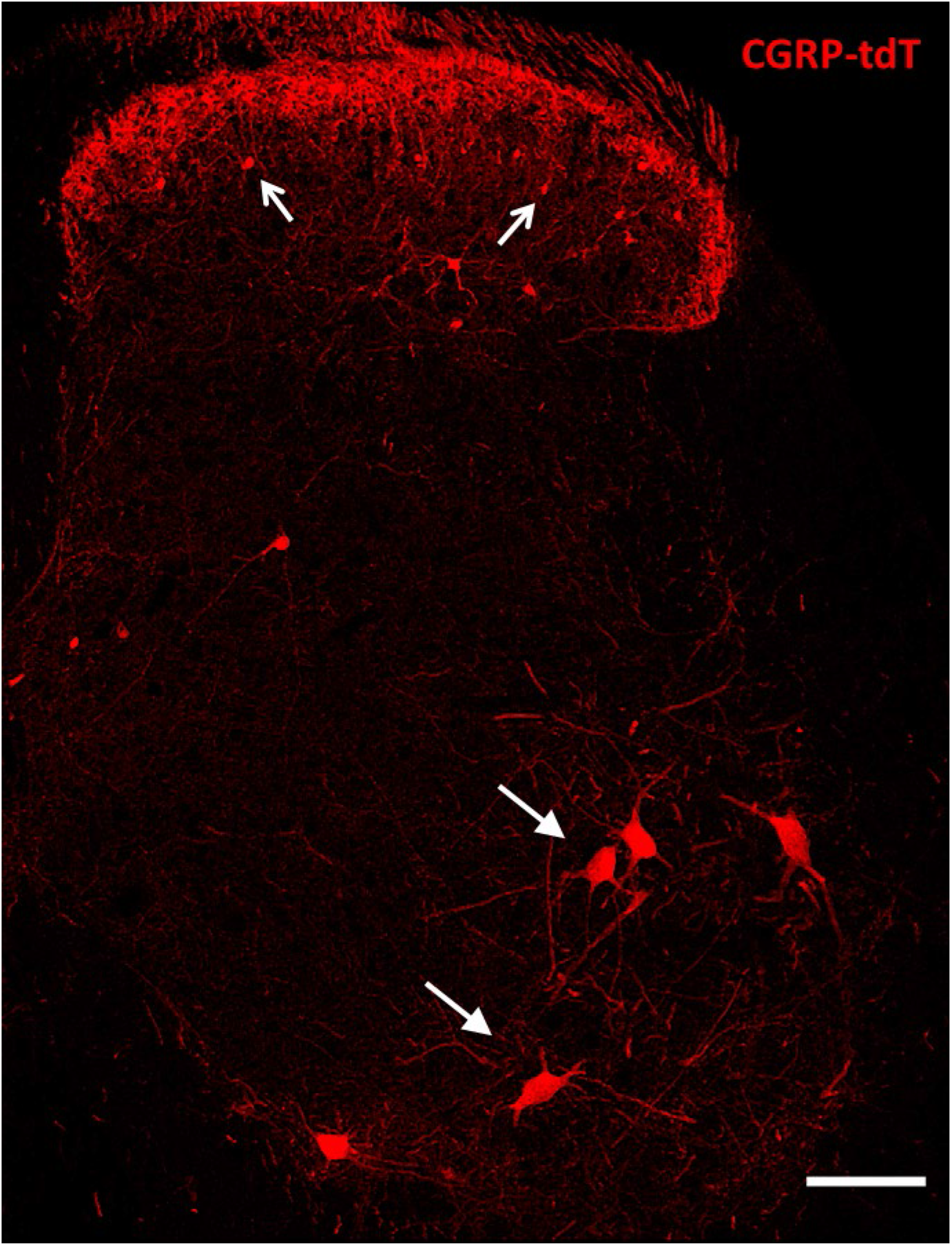
CGRP-tdTomato expression in the lumbar spinal cord. Lumbar section from a CGRP-tdTomato mouse. Arrows point to intensely fluorescent CGRP-tdTomato neurons in lamina III of the dorsal horn and ventral horn motoneurons. Laminae I and II (substantia gelatinosa) contain a dense array of fluorescent processes originating from CGRP-expressing primary sensory neurons. Scale bars: 100 μm.

**Supplementary Figure 2.**
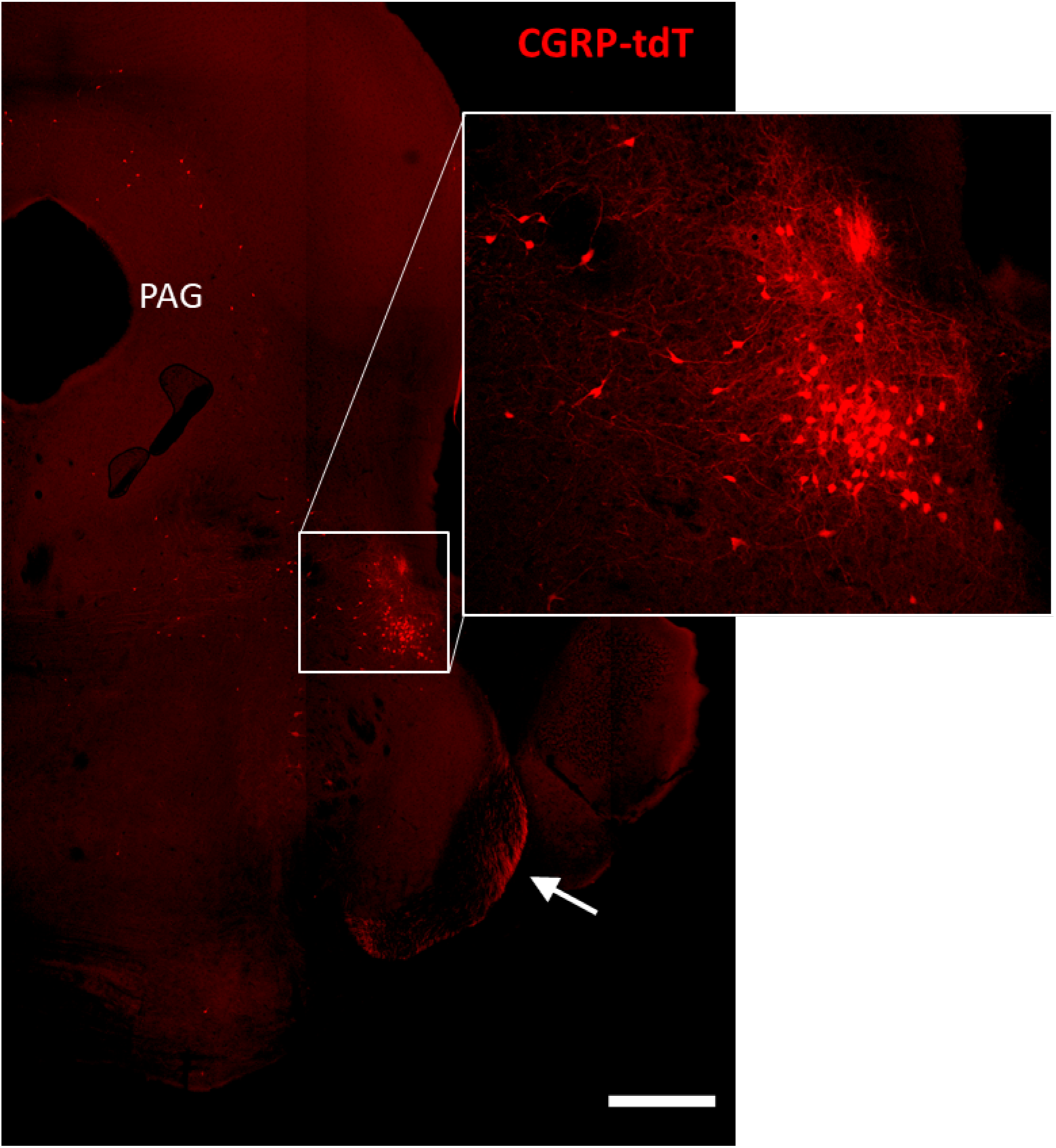
CGRP-tdTomato expression in the parabrachial nucleus. CGRP-tdTomato fluorescence at a caudal midbrain/rostral pontine level. The boxed area in the main figure shows intensely labelled neurons in the external lateral parabrachial nucleus, which are shown at higher magnification in the inset. The periaqueductal gray (PAG) also contains scattered CGRP-tdTomato-expressing neurons. The arrow points to primary afferent axons originating from the trigeminal ganglion. Scale bar: 500 μm.

**Supplementary Figure 3.**
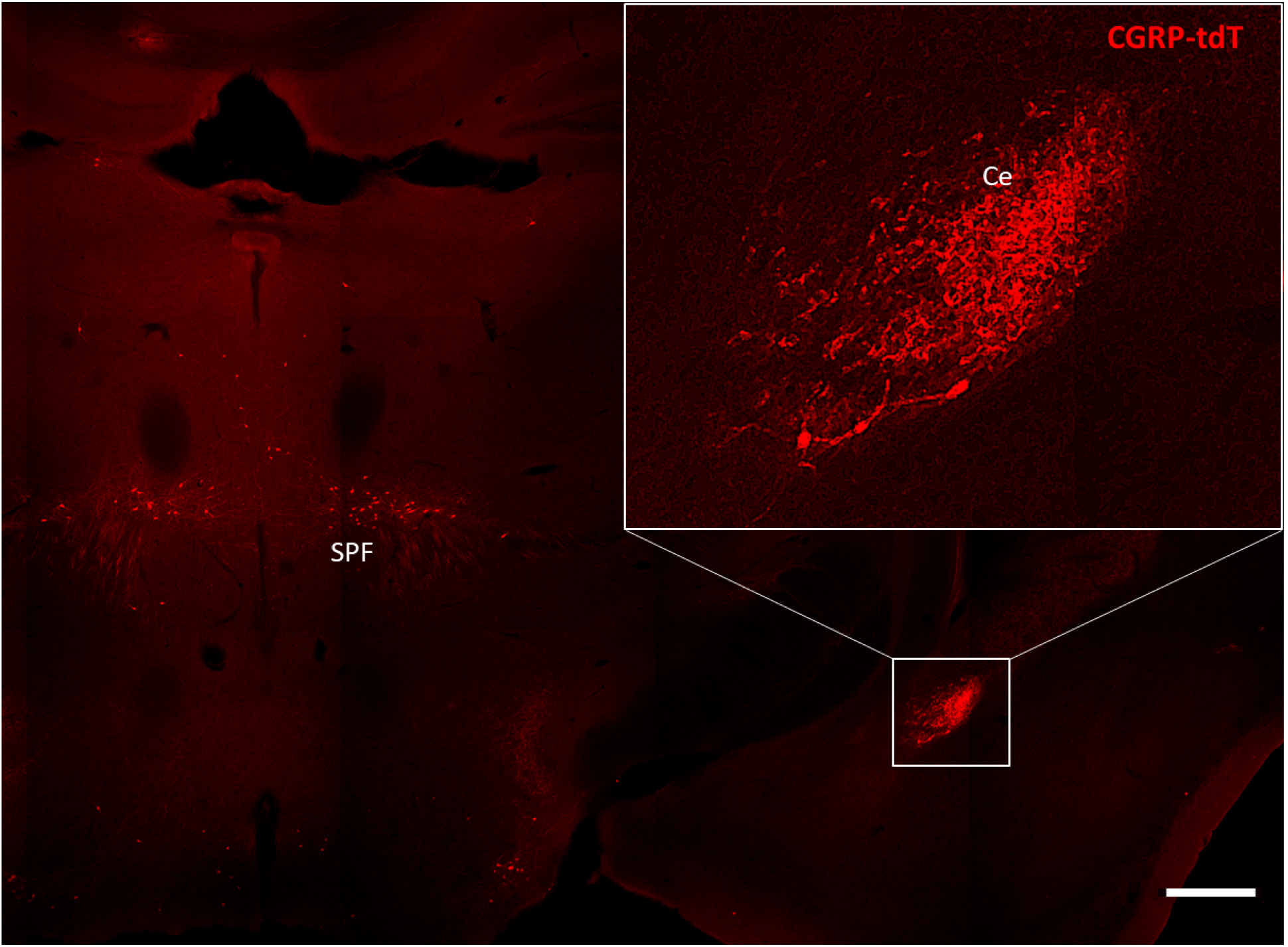
CGRP-tdTomato expression in the amygdala. CGRP-tdTomato fluorescence at a thalamic level. The boxed area in the main figure shows the dense plexus of CGRP-tdTomato fluorescent processes in the central nucleus of the amygdala (Ce). The section also illustrates labelled neurons in the subparafascicular nucleus of the thalamus (SPF). Scale bar: 500 μm.

**Supplementary Figure 4.**
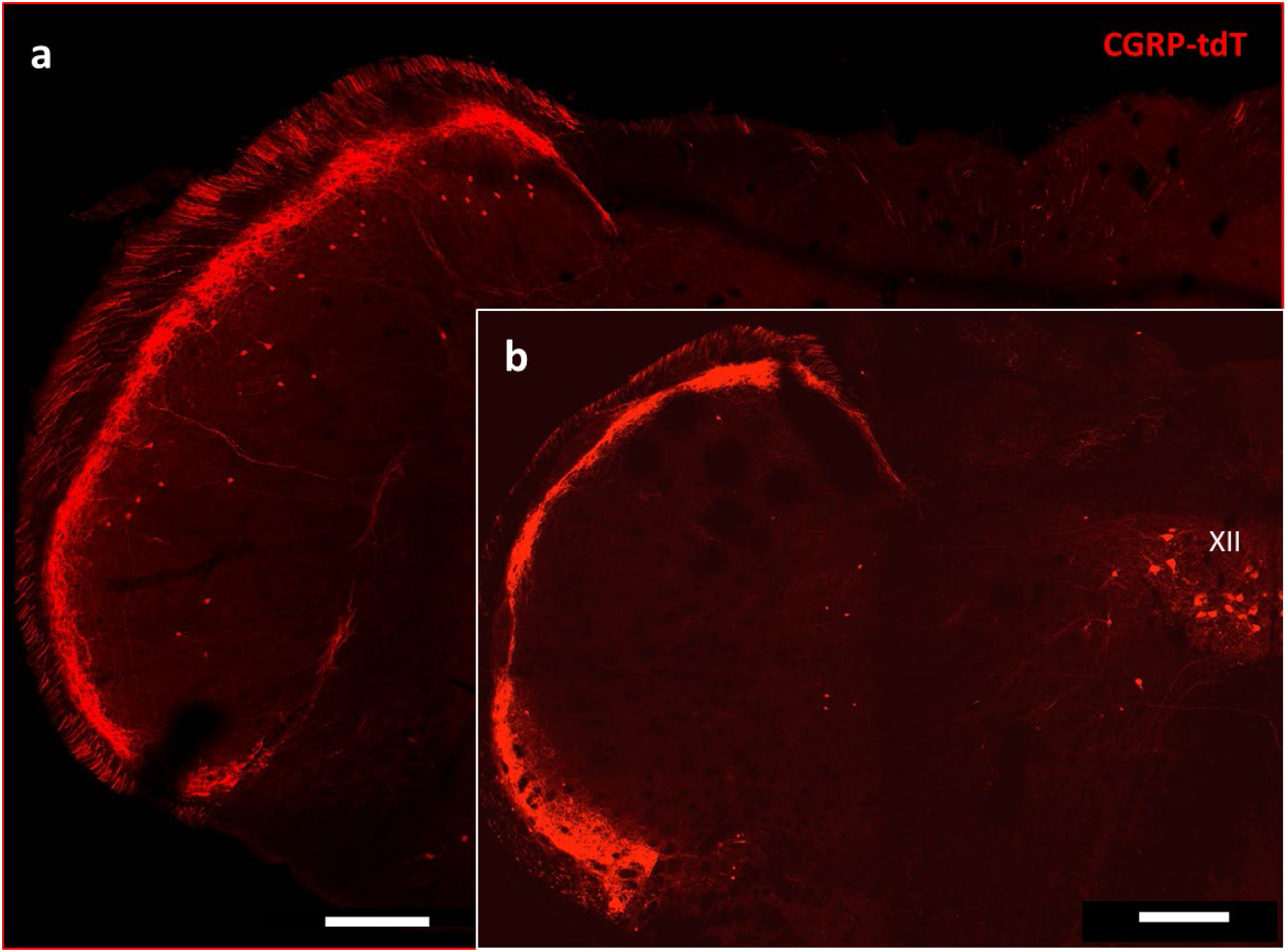
CGRP-tdTomato expression in the trigeminal nucleus caudalis. Caudal levels of the medulla (a) contain greater numbers of CGRP-tdTomato neurons in lamina III of the nucleus caudalis compared to more rostral levels (b). The latter level includes CGRP-tdTomato-expressing motoneurons in the hypoglossal nucleus (XII), Scale bars: 500 μm.

**Supplementary Figure 5.**
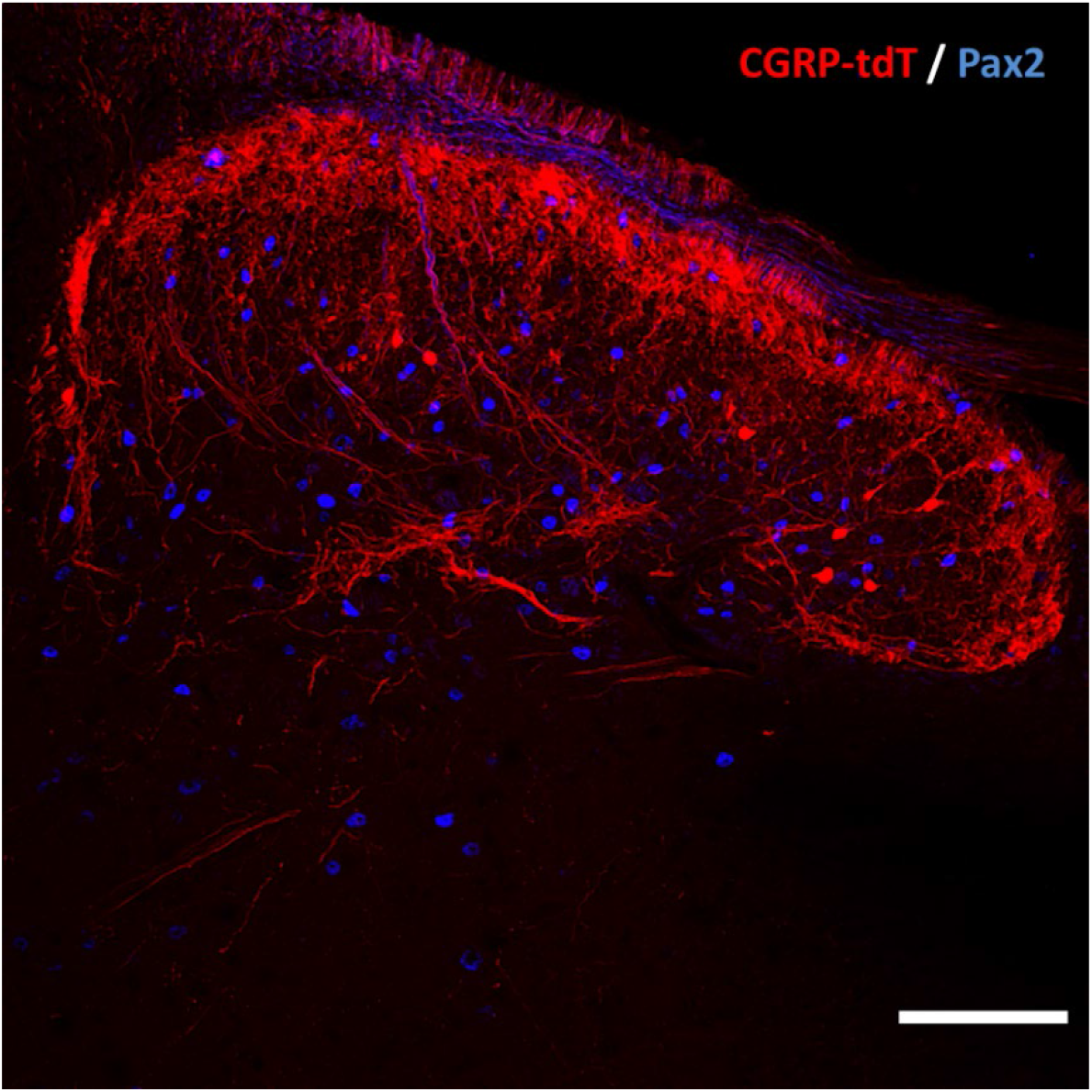
CGRP tdTomato interneurons are Pax2-negative. In lumbar dorsal horn, the absence of double labelling for CGRP-tdTomato (red) and Pax2 (blue)-immunoreactivity, a marker of inhibitory interneurons, indicates that the dorsal horn CGRP-tdTomato neurons are excitatory. Scale bar: 100 μm.

**Supplementary Figure 6:**
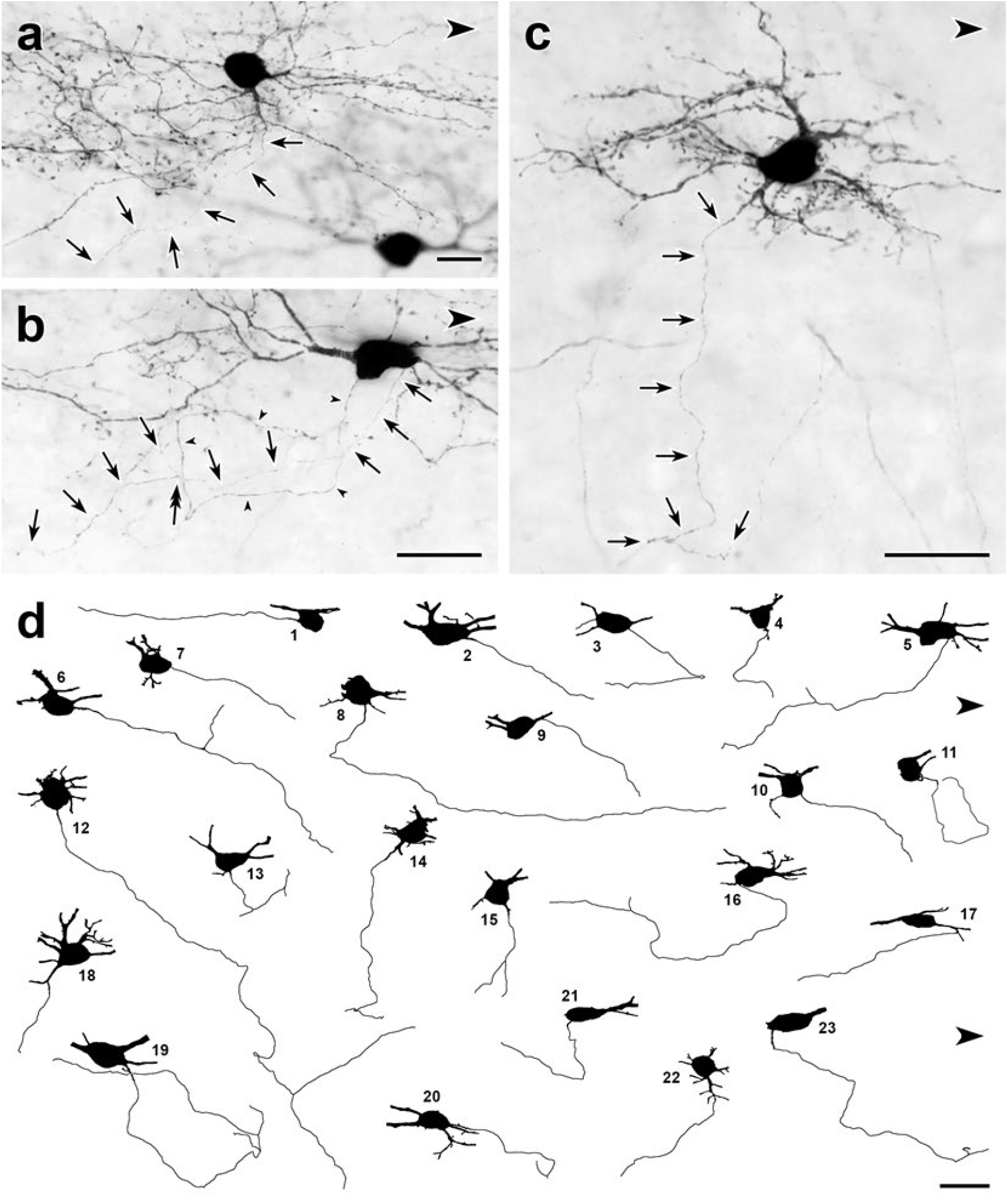
Dorsal horn CGRP interneurons have ventrally-directed axons. (**a-c**) Parasagittal 50 μm-thick sections of lumbar enlargement from CGRP-CreER-tdT mice that received intrathecal capsaicin treatment to reduce tdTomato-immunoreactivity from primary afferents. In most cases, the axons (arrows) of the dorsal horn CGRP-tdTomato interneurons travel ventrally. **a**, an axon arises from a ventrally-projecting primary dendrite, rather than the cell body. **b,** The axon (arrows) of this tdT-positive neuron arises from the caudal ventral surface of the cell body and travels ventrally and rostrally. The cell body also emits a very fine dendritic process, defined by the presence of spines (double-headed arrow). **c**) This heavily spine-laden, multipolar CGRP-tdTomato interneuron emits a ventrally directed axon from one of its dendrites. **d**, Drawings of 23 dorsal horn CGRP-tdTomato interneurons whose axons could be identified and traced. Each drawing shows the neuronal cell body and its axon as well as initial portions of its major primary dendrites. Nineteen of the axons originate from the ventral region of the cell body; 3 (**d14**, **d18**, **d22**), from a ventrally-projecting primary dendrite and one (**d17**) from a secondary dendrite close to its branch point off a primary dendrite. Most of the axons travel ventrally and caudally; some travel rostrally (e.g., **d1**, **d5**) and an occasional axon courses directly ventral (e.g., **d15**). After initially travelling ventro-caudally, 2 of the axons (**d11**, **d19**) looped dorsally and then began to travel rostrally. Eight of the axons bifurcated (**d5**, **d6**, **d12**, **d13**, **d14**, **d15**, **d16**, **d19**), all within 120 μm of their origin from a cell body. Scale bars: 20 μm.

**Supplementary Figure 7.**
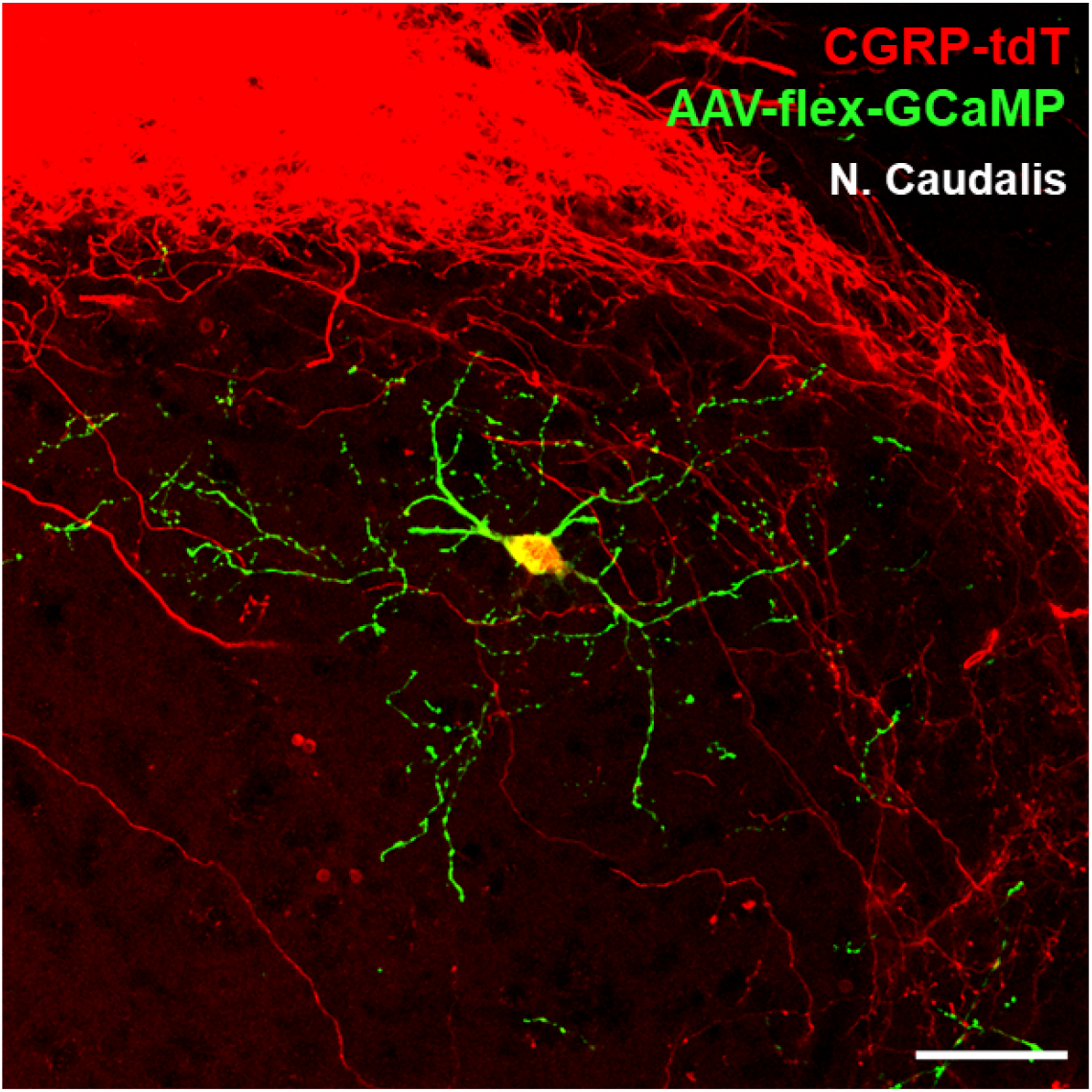
Radial morphology of the CGRP-tdTomato interneurons revealed after AAV injection. A CGRP-tdTomato neuron after injection of Cre-dependent AAV1-GCaMP6 (green) into the nucleus caudalis. This approach revealed a comparable morphology of the CGRP-tdTomato interneuron to that illustrated in Figure 4 and Supplementary Figures 6 a-c, but did not reveal distant axonal projections. Scale bar: 50 μm.

**Supplementary Figure 8.**
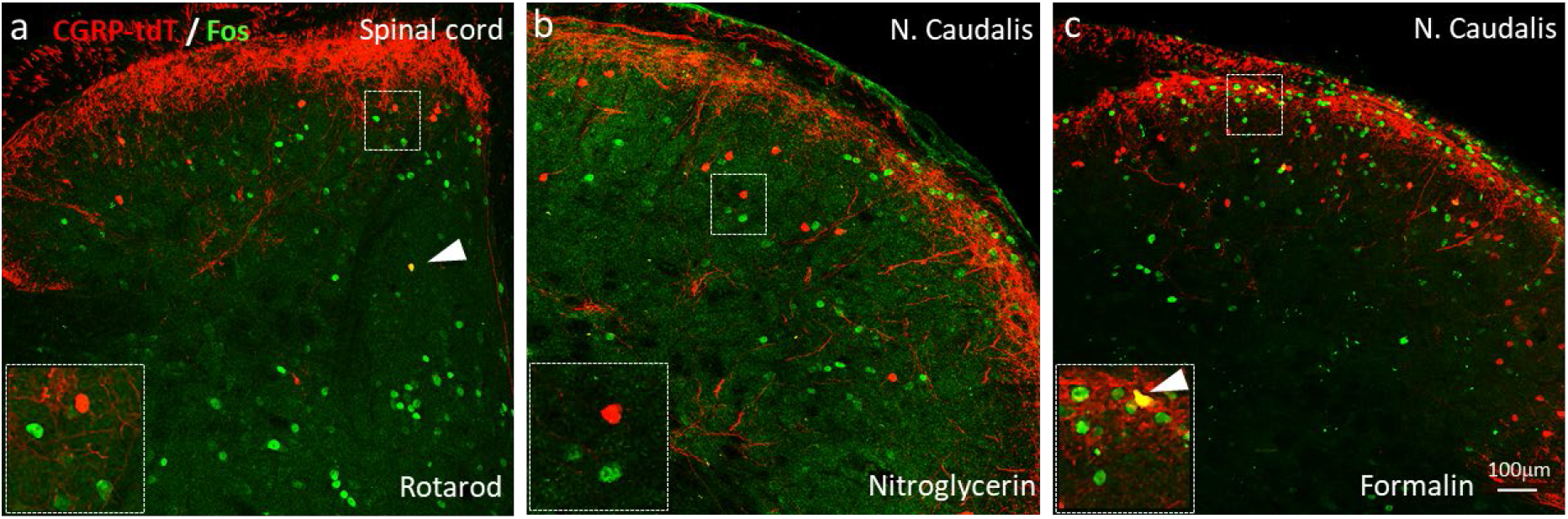
Neither noxious nor innocuous stimuli induce Fos expression in CGRP-tdTomato interneurons in control mice. **a**, Fos expression-immunoreactivity (green) in neurons of the lumbar dorsal horn after walking on a rotarod (90 mins). **b**. Fos-immunoreactivity in the neurons of the nucleus caudalis after systemic nitroglycerin injection (10 mg/kg, i.p.) or after a 2% formalin (10 μl) injection into the cheek **(c)** in unanesthetized mice. Insets illustrate higher magnification images of separate populations of Fos-immunoreactive and CGRP-tdTomato interneurons. Arrows in **a** and **c** point to rare double-labelled cells outside lamina III. Scale bar: 100 μm.

**Supplementary Figure 9.**
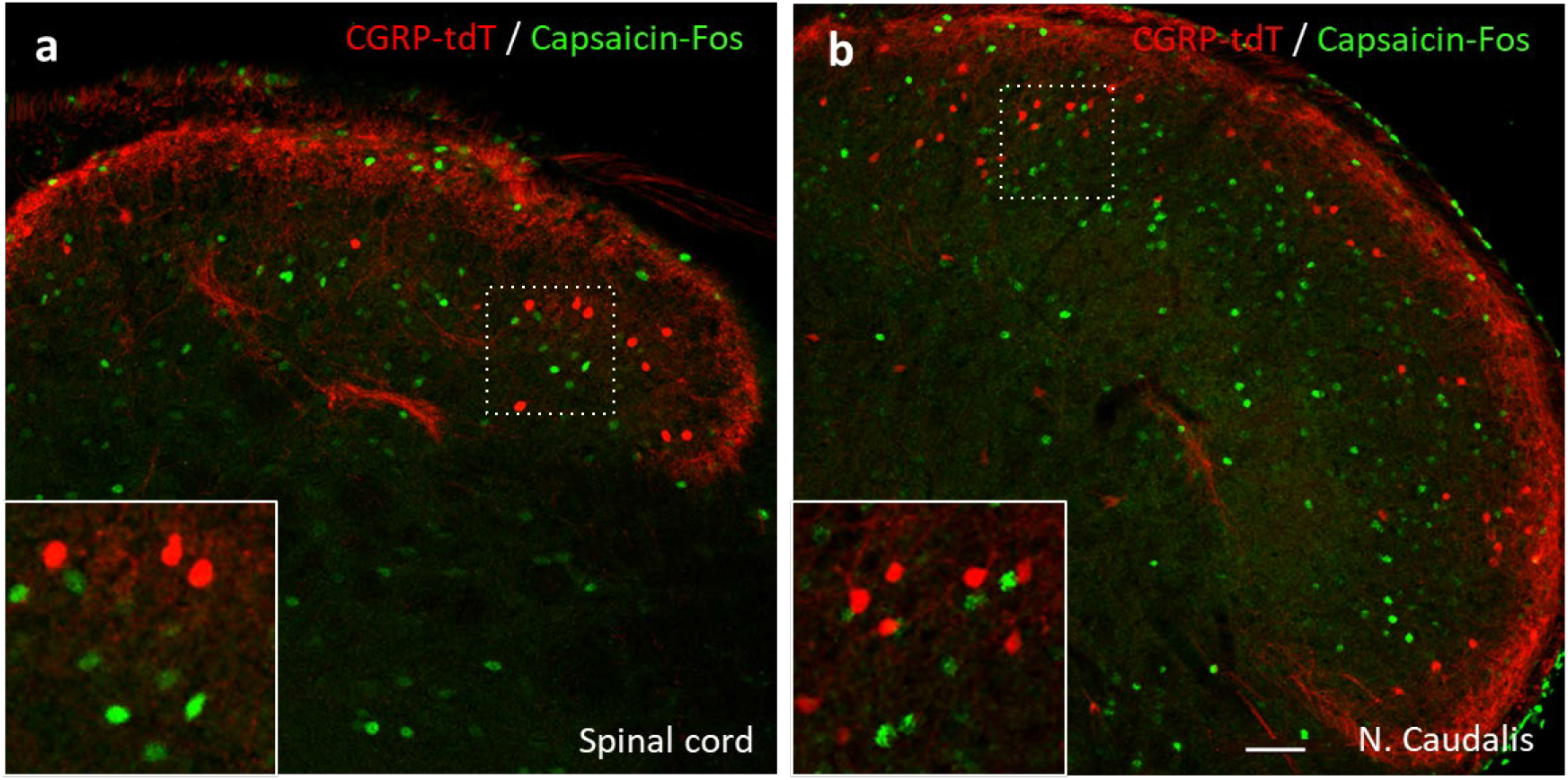
Capsaicin does not activate CGRP-tdTomato interneurons in the lumbar spinal cord or trigeminal nucleus caudalis. In anesthetized mice (2% isoflurane), a unilateral injection of 20 μl of capsaicin (1.0 μg/μl) into the hindpaw **(a)** or the cheek **(b)** did not induce Fos expression (green) in CGRP-tdTomato interneurons in the dorsal horn of the lumbar spinal cord (**a**) or in the nucleus caudalis (**b**). Insets show higher magnification views of boxed areas. Scale bar: 50 μm.

**Supplementary Figure 10.**
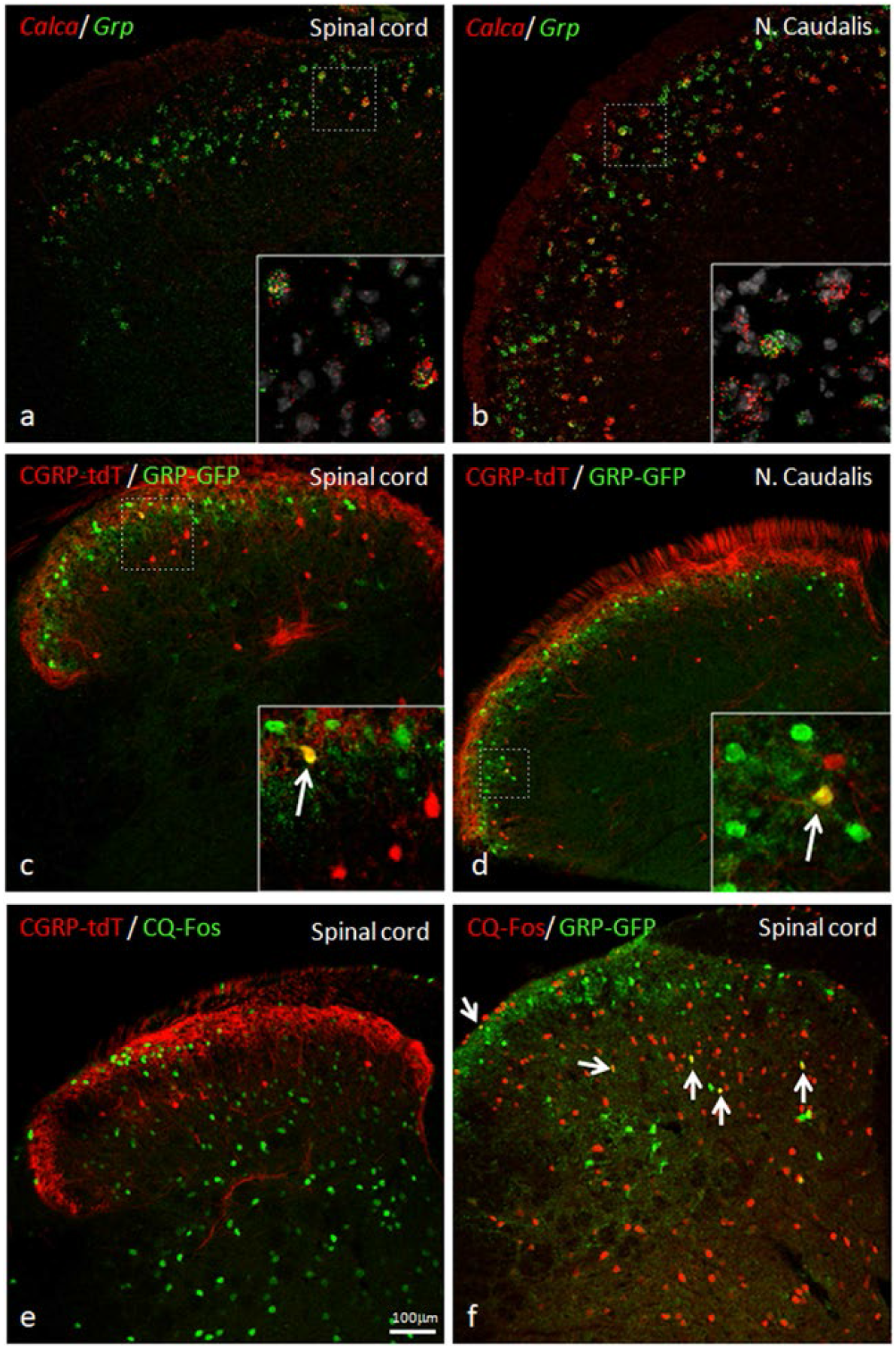
GRP, CGRP and pruritogen-evoked Fos expression. Double *in situ* hybridization for tdTomato (red) and GRP (green) illustrates considerable mRNA co-expression in neurons of the dorsal horn **(a)** and nucleus caudalis **(b)** of CGRP-tdTomato mice. In contrast, immunocytochemical localization of GRP and tdTomato in a tamoxifen-treated tdTomato-CGRP-CreER mouse that was crossed with a GRP-GFP reporter mouse revealed only occasional double labeling (arrow in inset) in the dorsal horn **(c)** or nucleus caudalis **(d).** Consistent with this minimal overlap, Fos expression in tdTomato-labeled CGRP interneurons was rare in response to a hindpaw injection of chloroquine (CQ; **e**). In contrast, many GRP-GFP interneurons were immunostained for Fos in response to CQ (arrows in **f**). As the mice were anesthetized the CQ-induced Fos was scratching-independent. Scale bar: 100 μm.

**Supplementary Table 1.** Electrophysiological properties of CGRP-tdTomato interneurons in the dorsal horn and nucleus caudalis. Most CGRP-tdTomato neurons showed delayed firing patterns (delayed 19, tonic 1, reluctant 2, single 2, no response 3). Based on electrical stimulation of dorsal roots, we conclude that CGRP interneurons in the lumbar cord predominantly receive monosynaptic input from Aβ primary afferent fibers.

| Membrane properties | Spinal cord | Caudalis |
| --- | --- | --- |
|  | 22 cells, 8 mice | 5 cells, 2 mice |
| Vm (mV) | -78.9 +/- 7.7 | -76 +/- 2.8 |
| Rheobase (pA) | 62.7 +/- 40.8 | 50.52 +/- 26.13 |
| AP thresh. | -41.3 +/- 15.2 | -48.15 +/- 9.44 |
| Cm (pF) | 40.3 +/- 14.7 | 44.55 +/- 12.63 |
| Rm (mOhm) | 603.1 +/- 296.8 | 710 +/- 463 |

| Firing pattern |  |  |
| --- | --- | --- |
| Delayed | 17/22 | 2/5 |
| Tonic | 1/22 | 0 |
| Reluctant | 1/22 | 1/5 |
| Single | 0 | 2/5 |
| No response | 3 | 0 |

**Supplementary Table 1.** Most CGRP-tdTomato neurons showed delayed firing patterns (delayed 19, tonic 1, reluctant 2, single 2, no response 3).
| Afferent input | 5 cells, 3 mice | n/a |
| --- | --- | --- |
| A-mono | 5/5 | n/a |
| C-poly & mono | 2/5 | n/a |

## REFERENCES

Abraira, V. E., Kuehn, E. D., Chirila, A. M., Springel, M. W., Toliver, A. A., Zimmerman, A. L., … Ginty, D. D. (2017). The cellular and synaptic architecture of the mechanosensory dorsal horn. Cell, 168(1-2), 295–310. doi:10.1016/j.cell.2016.12.010

Albisetti, G. W., Pagani, M., Platonova, E., Hosli, L., Johannssen, H. C., Fritschy, J. M., … Zeilhofer, H. U. (2019). Dorsal horn gastrin-releasing peptide expressing neurons transmit spinal itch but not pain signals. J Neurosci, 39(12), 2238–2250. doi:10.1523/JNEUROSCI.2559-18.2019

Basbaum, A. I., Bautista, D. M., Scherrer, G., … Julius, D. (2009). Cellular and molecular mechanisms of pain. Cell, 139(2), 267–284. doi:10.1016/j.cell.2009.09.028

Bates, E. A., Nikai, T., Brennan, K. C., Fu, Y. H., Charles, A. C., Basbaum, A. I., … Ahn, A. H. (2010). Sumatriptan alleviates nitroglycerin-induced mechanical and thermal allodynia in mice. Cephalalgia, 30(2), 170–178. doi:10.1111/j.1468-2982.2009.01864.x

Beal, J. A., … Cooper, M. H. (1978). The neurons in the gelatinosal complex (Laminae II and III) of the monkey (Macaca mulatta): a Golgi study. J Comp Neurol, 179(1), 89–121. doi:10.1002/cne.901790107

Bourane, S., Duan, B., Koch, S. C., Dalet, A., Britz, O., Garcia-Campmany, L., … Goulding, M. (2015). Gate control of mechanical itch by a subpopulation of spinal cord interneurons. Science, 350(6260), 550–554. doi:10.1126/science.aac8653

Bourane, S., Grossmann, K. S., Britz, O., Dalet, A., Del Barrio, M. G., Stam, F. J., … Goulding, M. (2015). Identification of a spinal circuit for light touch and fine motor control. Cell, 160(3), 503–515. doi:10.1016/j.cell.2015.01.011

Boyle, K. A., Gradwell, M. A., Yasaka, T., Dickie, A. C., Polgar, E., Ganley, R. P., … Hughes, D. I. (2019). Defining a spinal microcircuit that gates myelinated afferent input: implications for tactile allodynia. Cell Rep, 28(2), 526–540. doi:10.1016/j.celrep.2019.06.040

Brain, S. D., Williams, T. J., Tippins, J. R., Morris, H. R., … MacIntyre, I. (1985). Calcitonin gene-related peptide is a potent vasodilator. Nature, 313(5997), 54–56. doi:10.1038/313054a0

Cavanaugh, D. J., Lee, H., Lo, L., Shields, S. D., Zylka, M. J., Basbaum, A. I., … Anderson, D. J. (2009). Distinct subsets of unmyelinated primary sensory fibers mediate behavioral responses to noxious thermal and mechanical stimuli. Proc Natl Acad Sci U S A, 106(22), 9075–9080. doi:10.1073/pnas.0901507106

Chaplan, S. R., Bach, F. W., Pogrel, J. W., Chung, J. M., … Yaksh, T. L. (1994). Quantitative assessment of tactile allodynia in the rat paw. J Neurosci Methods, 53(1), 55–63. doi:10.1016/0165-0270(94)90144-9

Cheng, L., Duan, B., Huang, T., Zhang, Y., Chen, Y., Britz, O., … Ma, Q. (2017). Identification of spinal circuits involved in touch-evoked dynamic mechanical pain. Nat Neurosci, 20(6), 804–814. doi:10.1038/nn.4549

Cowie, A. M., Moehring, F., O’Hara, C., … Stucky, C. L. (2018). Optogenetic inhibition of CGRPalpha sensory neurons reveals their distinct roles in neuropathic and incisional Pain. J Neurosci, 38(25), 5807–5825. doi:10.1523/JNEUROSCI.3565-17.2018

Dickie, A. C., Bell, A. M., Iwagaki, N., Polgar, E., Gutierrez-Mecinas, M., Kelly, R., … Todd, A. J. (2019). Morphological and functional properties distinguish the substance P and gastrin-releasing peptide subsets of excitatory interneuron in the spinal cord dorsal horn. Pain, 160(2), 442–462. doi:10.1097/j.pain.0000000000001406

Duan, B., Cheng, L., Bourane, S., Britz, O., Padilla, C., Garcia-Campmany, L., … Ma, Q. (2014). Identification of spinal circuits transmitting and gating mechanical pain. Cell, 159(6), 1417–1432. doi:10.1016/j.cell.2014.11.003

Etlin, A., Braz, J. M., Kuhn, J. A., Wang, X., Hamel, K. A., Llewellyn-Smith, I. J., … Basbaum, I. (2016). Functional synaptic integration of forebrain GABAergic precursors into the adult spinal cord. J Neurosci, 36(46), 11634–11645. doi:10.1523/JNEUROSCI.2301-16.2016

Francois, A., Low, S. A., Sypek, E. I., Christensen, A. J., Sotoudeh, C., Beier, K. T., … Scherrer, G. (2017). A brainstem-spinal cord inhibitory circuit for mechanical pain modulation by GABA and enkephalins. Neuron, 93(4), 822–839. doi:10.1016/j.neuron.2017.01.008

Grudt, T. J., … Perl, E. R. (2002). Correlations between neuronal morphology and electrophysiological features in the rodent superficial dorsal horn. J Physiol, 540(Pt 1), 189–207. doi:10.1113/jphysiol.2001.012890

Haring, M., Zeisel, A., Hochgerner, H., Rinwa, P., Jakobsson, J. E. T., Lonnerberg, P., … Ernfors, P. (2018). Neuronal atlas of the dorsal horn defines its architecture and links sensory input to transcriptional cell types. Nat Neurosci, 21(6), 869–880. doi:10.1038/s41593-018-0141-1

Ho, T. W., Edvinsson, L., … Goadsby, P. J. (2010). CGRP and its receptors provide new insights into migraine pathophysiology. Nat Rev Neurol, 6(10), 573–582. doi:10.1038/nrneurol.2010.127

Huang, T., Lin, S. H., Malewicz, N. M., Zhang, Y., Zhang, Y., Goulding, M., … Ma, Q. (2019). Identifying the pathways required for coping behaviours associated with sustained pain. Nature, 565(7737), 86–90. doi:10.1038/s41586-018-0793-8

Imlach, W. L., Bhola, R. F., Mohammadi, S. A., … Christie, M. J. (2016). Glycinergic dysfunction in a subpopulation of dorsal horn interneurons in a rat model of neuropathic pain. Sci Rep, 6, 37104. doi:10.1038/srep37104

Kruger, L., Sternini, C., Brecha, N. C., … Mantyh, P. W. (1988). Distribution of calcitonin gene-related peptide immunoreactivity in relation to the rat central somatosensory projection. J Comp Neurol, 273(2), 149–162. doi:10.1002/cne.902730203

Liu, Y., Latremoliere, A., Li, X., Zhang, Z., Chen, M., Wang, X., … He, Z. (2018). Touch and tactile neuropathic pain sensitivity are set by corticospinal projections. Nature, 561(7724), 547–550. doi:10.1038/s41586-018-0515-2

Llewellyn-Smith, I. J., Basbaum, A. I., … Braz, J. M. (2018). Long-term, dynamic synaptic reorganization after GABAergic precursor cell transplantation into adult mouse spinal cord. J Comp Neurol, 526(3), 480–495. doi:10.1002/cne.24346

Llewellyn-Smith, I. J., Dicarlo, S. E., Collins, H. L., … Keast, J. R. (2005). Enkephalin-immunoreactive interneurons extensively innervate sympathetic preganglionic neurons regulating the pelvic viscera. J Comp Neurol, 488(3), 278–289. doi:10.1002/cne.20552

Lu, Y., Dong, H., Gao, Y., Gong, Y., Ren, Y., Gu, N., … Xiong, L. (2013). A feed-forward spinal cord glycinergic neural circuit gates mechanical allodynia. J Clin Invest, 123(9), 4050–4062. doi:10.1172/JCI70026

Malmberg, A. B., Chen, C., Tonegawa, S., … Basbaum, A. I. (1997). Preserved acute pain and reduced neuropathic pain in mice lacking PKCgamma. Science, 278(5336), 279–283. doi:10.1126/science.278.5336.279

Maxwell, D. J. (1985). Combined light and electron microscopy of Golgi-labelled neurons in lamina III of the feline spinal cord. J Anat, 141, 155–169.

McCoy, E. S., Taylor-Blake, B., … Zylka, M. J. (2012). CGRPalpha-expressing sensory neurons respond to stimuli that evoke sensations of pain and itch. PLoS One, 7(5). doi:10.1371/journal.pone.0036355

Neumann, S., Braz, J. M., Skinner, K., Llewellyn-Smith, I. J., … Basbaum, A. I. (2008). Innocuous, not noxious, input activates PKCgamma interneurons of the spinal dorsal horn via myelinated afferent fibers. J Neurosci, 28(32), 7936–7944. doi:10.1523/JNEUROSCI.1259-08.2008

Oliveira, A. L., Hydling, F., Olsson, E., Shi, T., Edwards, R. H., Fujiyama, F., … Meister, B. (2003). Cellular localization of three vesicular glutamate transporter mRNAs and proteins in rat spinal cord and dorsal root ganglia. Synapse, 50(2), 117–129. doi:10.1002/syn.10249

Patil, M. J., Hovhannisyan, A. H., … Akopian, A. N. (2018). Characteristics of sensory neuronal groups in CGRP-cre-ER reporter mice: Comparison to Nav1.8-cre, TRPV1-cre and TRPV1-GFP mouse lines. PLoS One, 13(6), e0198601. doi:10.1371/journal.pone.0198601

Peirs, C., Williams, S. P., Zhao, X., Walsh, C. E., Gedeon, J. Y., Cagle, N. E., … Seal, R. P. (2015). Dorsal horn circuits for persistent mechanical pain. Neuron, 87(4), 797–812. doi:10.1016/j.neuron.2015.07.029

Petitjean, H., Bourojeni, F. B., Tsao, D., Davidova, A., Sotocinal, S. G., Mogil, J. S., … Sharif-Naeini, R. (2019). Recruitment of spinoparabrachial neurons by dorsal horn calretinin neurons. Cell Rep, 28(6), 1429–1438. doi:10.1016/j.celrep.2019.07.048

Petitjean, H., Pawlowski, S. A., Fraine, S. L., Sharif, B., Hamad, D., Fatima, T., … Sharif-Naeini, R. (2015). Dorsal horn parvalbumin neurons are gate-keepers of touch-evoked pain after nerve injury. Cell Rep, 13(6), 1246–1257. doi:10.1016/j.celrep.2015.09.080

Punnakkal, P., von Schoultz, C., Haenraets, K., Wildner, H., … Zeilhofer, H. U. (2014). Morphological, biophysical and synaptic properties of glutamatergic neurons of the mouse spinal dorsal horn. J Physiol, 592(4), 759–776. doi:10.1113/jphysiol.2013.264937

Ryu, P. D., Gerber, G., Murase, K., … Randic, M. (1988). Calcitonin gene-related peptide enhances calcium current of rat dorsal root ganglion neurons and spinal excitatory synaptic transmission. Neurosci Lett, 89(3), 305–312. doi:10.1016/0304-3940(88)90544-7

Sathyamurthy, A., Johnson, K. R., Matson, K. J. E., Dobrott, C. I., Li, L., Ryba, A. R., … Levine, A. J. (2018). Massively parallel single nucleus transcriptional profiling defines spinal cord neurons and their activity during behavior. Cell Rep, 22(8), 2216–2225. doi:10.1016/j.celrep.2018.02.003

Sherman, S. E., … Loomis, C. W. (1996). Strychnine-sensitive modulation is selective for non-noxious somatosensory input in the spinal cord of the rat. Pain, 66(2-3), 321–330. doi:10.1016/0304-3959(96)03063-1

Shields, S. D., Eckert, W. A., 3rd, … Basbaum, A. I. (2003). Spared nerve injury model of neuropathic pain in the mouse: a behavioral and anatomic analysis. J Pain, 4(8), 465–470. doi:10.1067/s1526-5900(03)00781-8

Smith, K. M., Browne, T. J., Davis, O. C., Coyle, A., Boyle, K. A., Watanabe, M., … Graham, A. (2019). Calretinin positive neurons form an excitatory amplifier network in the spinal cord dorsal horn. Elife, 8. doi:10.7554/eLife.49190

Solorzano, C., Villafuerte, D., Meda, K., Cevikbas, F., Braz, J., Sharif-Naeini, R., … Basbaum, A. I. (2015). Primary afferent and spinal cord expression of gastrin-releasing peptide: message, protein, and antibody concerns. J Neurosci, 35(2), 648–657. doi:10.1523/JNEUROSCI.2955-14.2015

Song, H., Yao, E., Lin, C., Gacayan, R., Chen, M. H., … Chuang, P. T. (2012). Functional characterization of pulmonary neuroendocrine cells in lung development, injury, and tumorigenesis. Proc Natl Acad Sci U S A, 109(43), 17531–17536. doi:10.1073/pnas.1207238109

Sun, Y. G., … Chen, Z. F. (2007). A gastrin-releasing peptide receptor mediates the itch sensation in the spinal cord. Nature, 448(7154), 700–703. Retrieved from https://www.ncbi.nlm.nih.gov/pubmed/17653196. doi:10.1038/nature06029

Tie-Jun, S. S., Xu, Z., … Hokfelt, T. (2001). The expression of calcitonin gene-related peptide in dorsal horn neurons of the mouse lumbar spinal cord. Neuroreport, 12(4), 739–743. doi:10.1097/00001756-200103260-00025

Torsney, C., … MacDermott, A. B. (2006). Disinhibition opens the gate to pathological pain signaling in superficial neurokinin 1 receptor-expressing neurons in rat spinal cord. J Neurosci, 26(6), 1833–1843. doi:10.1523/JNEUROSCI.4584-05.2006

Wercberger, R., … Basbaum, A.I. (2019). Spinal cord projection neurons: a superficial, and also deep analysis. Curr Opin Physiol, 11, 109–115. doi:10.1016/j.cophys.2019.10.002

Woolf, C., … Wiesenfeld-Hallin, Z. (1986). Substance P and calcitonin gene-related peptide synergistically modulate the gain of the nociceptive flexor withdrawal reflex in the rat. Neurosci Lett, 66(2), 226–230. doi:10.1016/0304-3940(86)90195-3

Yasaka, T., Tiong, S. Y., Hughes, D. I., Riddell, J. S., … Todd, A. J. (2010). Populations of inhibitory and excitatory interneurons in lamina II of the adult rat spinal dorsal horn revealed by a combined electrophysiological and anatomical approach. Pain, 151(2), 475–488. doi:10.1016/j.pain.2010.08.008

Yasui, Y., Saper, C. B., … Cechetto, D. F. (1991). Calcitonin gene-related peptide (CGRP) immunoreactive projections from the thalamus to the striatum and amygdala in the rat. J Comp Neurol, 308(2), 293–310. doi:10.1002/cne.903080212

Yoshida, A., Dostrovsky, J. O., Sessle, B. J., … Chiang, C. Y. (1991). Trigeminal projections to the nucleus submedius of the thalamus in the rat. J Comp Neurol, 307(4), 609–625. doi:10.1002/cne.903070408

